# LDpred-funct: incorporating functional priors improves polygenic prediction accuracy in UK Biobank and 23andMe data sets

**DOI:** 10.1101/375337

**Authors:** Carla Márquez-Luna, Steven Gazal, Po-Ru Loh, Samuel S. Kim, Nicholas Furlotte, Adam Auton, 23andMe Research Team, Alkes L. Price

## Abstract

Genetic variants in functional regions of the genome are enriched for complex trait heritability. Here, we introduce a new method for polygenic prediction, LDpred-funct, that leverages trait-specific functional priors to increase prediction accuracy. We fit priors using the recently developed baseline-LD model, which includes coding, conserved, regulatory and LD-related annotations. We analytically estimate posterior mean causal effect sizes and then use cross-validation to regularize these estimates, improving prediction accuracy for sparse architectures. LDpred-funct attained higher prediction accuracy than other polygenic prediction methods in simulations using real genotypes. We applied LDpred-funct to predict 21 highly heritable traits in the UK Biobank. We used association statistics from British-ancestry samples as training data (avg *N*=373K) and samples of other European ancestries as validation data (avg *N*=22K), to minimize confounding. LDpred-funct attained a +4.6% relative improvement in average prediction accuracy (avg prediction *R*^2^=0.144; highest *R*^2^=0.413 for height) compared to SBayesR (the best method that does not incorporate functional information). For height, meta-analyzing training data from UK Biobank and 23andMe cohorts (total *N*=1107K; higher heritability in UK Biobank cohort) increased prediction *R*^2^ to 0.431. Our results show that incorporating functional priors improves polygenic prediction accuracy, consistent with the functional architecture of complex traits.

## Introduction

Genetic variants in functional regions of the genome are enriched for complex trait heritability^1–6^. In this study, we aim to leverage functional priors to improve polygenic prediction^7,8^. Several studies have shown that incorporating prior distributions on causal effect sizes can improve prediction accuracy^9–16^, compared to standard Best Linear Unbiased Prediction (BLUP) or Pruning+Thresholding methods^17–22^. Recent efforts to incorporate functional information have produced promising results^23,24^ (see P+T-funct-LASSO and AnnoPred results in all main figures below), but may be limited by dichotomizing between functional and non-functional variants^23^ or restricting their analyses to genotyped variants^24^.

Here, we introduce a new method, LDpred-funct, for leveraging trait-specific functional priors to increase polygenic prediction accuracy. We fit functional priors using our recently developed baseline-LD model^25^, which includes coding, conserved, regulatory and LD-related annotations. LDpred-funct first analytically estimates posterior mean causal effect sizes, accounting for functional priors and LD between variants. LDpred-funct then uses cross-validation within validation samples to regularize causal effect size estimates in bins of different magnitude, improving prediction accuracy for sparse architectures. We show that LDpred-funct attains higher polygenic prediction accuracy than other methods in simulations with real genotypes, analyses of 21 highly heritable UK Biobank traits, and meta-analyses of height using training data from UK Biobank and 23andMe cohorts.

## Methods

### Polygenic prediction methods

We compared 7 main prediction methods: Pruning+Thresholding^18,19^ (P+T), LDpred^16^, SBayesR^9^, P+T with functionally informed LASSO shrinkage^23^ (P+T-funct-LASSO), AnnoPred^24^, our new LDpred-funct-inf method, and our new LDpred-funct method; we also included LDpred-inf^16^, which is known to attain lower prediction accuracy than LDpred^16^, in some of our secondary analyses. P+T, LDpred-inf, LDpred and SBayesR are polygenic prediction methods that do not use functional annotations; we did not include the recently developed RSS^12^ and SBLUP^11^ methods in our comparisons, because ref. 9 reported that SBayesR performed as well or better than both RSS and SBLUP and was more computationally efficient (Figure 2 and Figure S18 of ref. 9). P+T-funct-LASSO is a modification of P+T that corrects marginal effect sizes for winner’s curse, accounting for functional annotations. AnnoPred is which uses a Bayesian framework to incorporate functional annotations. LDpred-funct-inf is an improvement of LDpred-inf that incorporates functionally informed priors on causal effect sizes. LDpred-funct is an improvement of LDpred-funct-inf that uses cross-validation to regularize posterior mean causal effect size estimates, improving prediction accuracy for sparse architectures. Each method is described in greater detail below. In both simulations and analyses of real traits, we used squared correlation (*R*^2^) between predicted phenotype and true phenotype in a held-out set of samples as our primary measure of prediction accuracy.

#### P+T

The P+T method builds a polygenic risk score (PRS) using a subset of independent SNPs obtained via informed LD-pruning^19^ (also known as LD-clumping) followed by P-value thresholding^18^. Specifically, the method has two parameters, 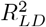 and *P_T_*, and proceeds as follows. First, the method prunes SNPs based on a pairwise threshold 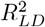, removing the less significant SNP in each pair. Second, the method restricts to SNPs with an association P-value below the significance threshold *P_T_*. Letting *M* be the number of SNPs remaining after LD-clumping, polygenic risk scores (PRS) are computed as

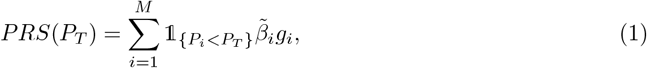

where 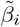 are normalized marginal effect size estimates and *g_i_* is a vector of normalized genotypes for SNP *i*. The parameters 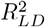 and *P_T_* are commonly tuned using validation data to optimize prediction accuracy^18,19^. While in theory this procedure is susceptible to overfitting, in practice, validation sample sizes are typically large, and 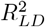 and *P_T_* are selected from a small discrete set of parameter choices, so that overfitting is considered to have a negligible effectM^8,19,26^. Accordingly, in this work, we consider 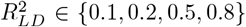 and *P_T_* ∈ {1, 0.3, 0.1,0.03, 0.01, 0.003, 0.001, 3 * 10^−4^, 10^−4^, 3 * 10^−5^, 10^−5^,10^−6^, 10^−7^,10^−8^}, and we always report results corresponding to the best choices of these parameters. The P+T method is implemented in the PLINK software (see Web Resources).

#### LDpred-inf

The LDpred-inf method estimates posterior mean causal effect sizes under an infinitesimal model, accounting for LD^16^. The infinitesimal model assumes that normalized causal effect sizes have prior distribution *β_i_* ~ *N* (0, *σ*^2^), where 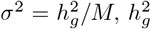 is the SNP-heritability, and *M* is the number of SNPs. The posterior mean causal effect sizes are

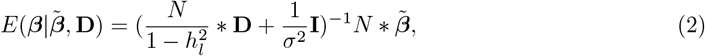

where **D** is the LD matrix between markers, **I** is the identity matrix, *N* is the training sample size, 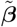 is the vector of marginal association statistics, and 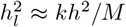 is the heritability of the *k* SNPs in the region of LD; following ref. 16 we use the approximation 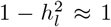, which is appropriate when *M* >> *k*. D is typically estimated using validation data, restricting to non-overlapping LD windows. We used the default LD window size, which is M/3000. 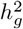 can be estimated from raw genotype/phenotype data^27,28^ (the approach that we use here; see below), or can be estimated from summary statistics using the aggregate estimator as described in ref. 16. To approximate the normalized marginal effect size ref. 16 uses the p-values to obtain absolute Z scores and then multiplies absolute Z scores by the sign of the estimated effect size. When sample sizes are very large, p-values may be rounded to zero, in which case we approximate normalized marginal effect sizes 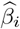 by 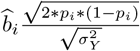, where 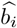 is the per-allele marginal effect size estimate, *p_i_* is the minor allele frequency of SNP *i*, and 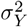 is the phenotypic variance in the training data. This applies to all the methods that use normalized effect sizes. Although the published version of LDpred requires a matrix inversion (Equation 2), we have implemented a computational speedup that computes the posterior mean causal effect sizes by efficiently solving^29^ the system of linear equations 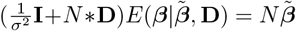.

#### LDpred

The LDpred method is an extension of LDpred-inf that uses a point-normal prior to estimate posterior mean effect sizes via Markov Chain Monte Carlo (MCMC)^16^. It assumes a Gaussian mixture prior: 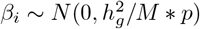 with probability *p*, and *β_i_* ~ 0 with probability 1 − *p*, where *p* is the proportion of causal SNPs. The method is optimized by considering different values of *p* (1E-4, 3E-4, 1E-3, 3E-3, 0.01,0.03,0.1,0.3,1); in the special case where 100% of SNPs are assumed to be causal, LDpred is roughly equivalent to LDpred-inf. We excluded SNPs from long-range LD regions (reported in ref. 30), as our secondary analyses showed that including these regions was suboptimal, consistent with ref. 9.

#### SBayesR

The SBayesR method infers posterior mean causal effect sizes from GWAS summary statistics and an LD matrix^9^. It assumes a finite mixture of normal distributions to account for sparsity, defined as: 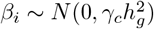 with probability *π_c_*, where *c* ranges from 1 to *C*, the total number of components in the mixture model. We used as input the recommended parameters from ref. 9, with *C* = 4 mixtures with parameters γ*_c_* = (0, 0.01, 0.1, 1.0). The method requires a shrunk LD matrix^12^. The authors of ref. 9 made available shrunk LD matrices estimated from 50,000 randomly selected white British individuals from the UK Biobank^30^ for two different SNPs sets. The 1.1M SNP set consists of 1,094,841 variants, constructed by restricting 1,365,446 SNPs from HapMap3^31^ to MAF > 0.01 and removing strand ambiguous SNPs and long-range LD regions (as reported in ref. 30). The 2.9M SNP set consists of 2,865,810 variants, constructed by applying LD-pruning (*R*^2^ > 0.99) to a larger set of 8 million variants from the UK Biobank^30^ with MAF > 0.01, overlapped with a previous large GWAS^32^ and present in 1000 Genomes^33^. We note that we could not scale the SBayesR analysis to the full set of 6,334,603 variants used in other analyses due to computational constraints. We used the 1.1M SNP set in our primary analyses as it achieved the highest average prediction *R*^2^ in our real traits analyses (see Results section), but we also considered the 2.9M SNP set in secondary analyses. For analyses that use BOLT-LMM summary statistics we used *N_effective_* as reported in ref. 27.

#### P+T-funct-LASSO

Ref. 23 proposed an extension of P+T that corrects the marginal effect sizes of SNPs for winner’s curse and incorporates external functional annotation data (P+T-funct-LASSO). The winner’s curse correction is performed by applying a LASSO shrinkage to the marginal association statistics of the PRS:

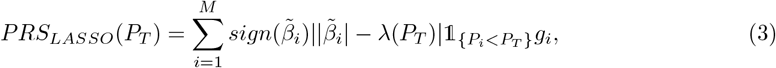

where 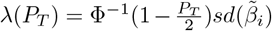, where Φ^−1^ is the inverse standard normal CDF. Functional annotations are incorporated via two disjoint SNPs sets, representing “high-prior” SNPs (HP) and “low-prior” SNPs (LP), respectively. We define the HP SNP set for P+T-funct-LASSO as the set of SNPs in the top 10% of expected per-SNP heritability under the baseline-LD model^25^, which includes coding, conserved, regulatory and LD-related annotations, whose enrichments are jointly estimated using stratified LD score regression^5,25^ (see Baseline-LD model annotations section). We also performed secondary analyses using the top 5% (P+T-funct-LASSO-top5%). We define *PRS_LASSO,HP_*(*P_HP_*) to be the PRS restricted to the HP SNP set, and *PRS_LASSO,LP_*(*P_LP_*) to be the PRS restricted to the LP SNP set, where *P_HP_* and *P_LP_* are the optimal significance thresholds for the HP and LP SNP sets, respectively. We define *PRS_LASSO_*(*P_HP_,P_LP_*) = *PRS_LASSO,HP_*(*P_HP_*)+*PRS_LASSO,LP_*(*P_LP_*). We also performed secondary analyses were we allow an additional regularization to the two PRS: *PRS_LASSO_*(*P_HP_,P_LP_*) = *α*_1_*PRS_LASSO,HP_*(*P_HP_*) + *α*_2_*PRS_LASSO,LP_*(*P_LP_*); we refer to this method as P+T-funct-LASSO-weighted.

#### AnnoPred

AnnoPred^24^ uses a Bayesian framework to incorporate functional priors while accounting for LD, optimizing prediction *R*^2^ over different assumed values of the proportion of causal SNPs. Ref. 24 proposed two different priors for use with AnnoPred. The first prior assumes the same proportion of causal SNPs but different causal effect size variance across functional annotations, and uses a point-normal prior to estimate posterior mean effect sizes via Markov Chain Monte Carlo (MCMC). In the special case where 100% of SNPs are assumed to be causal, AnnoPred is roughly equivalent to LDpred-funct-inf (see below). The second prior assumes different proportions of causal SNPs but the same causal effect size variance across functional annotations. We only consider the first prior, since the second prior cannot be extended to incorporate continuous-valued annotations from the baseline-LD model. We excluded SNPs from long-range LD regions (as reported in ref. 30) when running AnnoPred. We used the default LD window size, which is M/3000.

#### LDpred-funct-inf

We modify LDpred-inf to incorporate functionally informed priors on causal effect sizes using the baseline-LD model^25^, which includes coding, conserved, regulatory and LD-related annotations, whose enrichments are jointly estimated using stratified LD score regression^5,25^. Specifically, we assume that normalized causal effect sizes have prior distribution 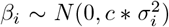, where 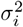 is the expected per-SNP heritability under the baseline-LD model (fit using training data only) and *c* is a normalizing constant such that 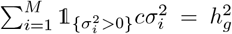, SNPs with 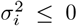 are removed, which is equivalent to setting 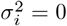. The posterior mean causal effect sizes are

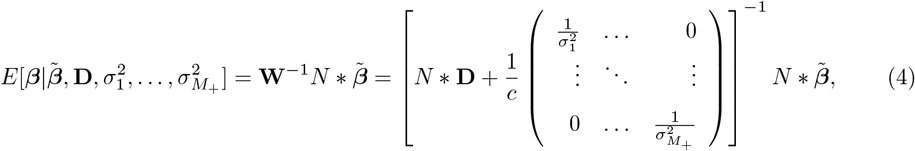

where *M*_+_ is the number of SNPs with 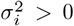. The posterior mean causal effect sizes are computed by solving the system of linear equations 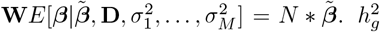 is estimated as described above (see LDpred-inf). **D** is estimated using validation data, restricting to windows of size 0.15%M_+_. In principle, it is possible to use banding to define the LD matrices, where LD between distant pair of SNPs (10 Mb or more) is rounded to zero^34^, but we elected to use the simpler window-based approach (as in ref. 16).

#### LDpred-funct

We modify LDpred-funct-inf to regularize posterior mean causal effect sizes using cross-validation. We rank the SNPs by their (absolute) posterior mean causal effect sizes, partition the SNPs into *K* bins (analogous to ref. 35) where each bin has roughly the same sum of squared posterior mean effect sizes, and determine the relative weights of each bin based on predictive value in the validation data. Intuitively if a bin is dominated by non-causal SNPs, the inferred relative weight will be lower than for a bin with a high proportion of causal SNPs. This non-parametric shrinkage approach can optimize prediction accuracy regardless of the genetic architecture. In detail, let 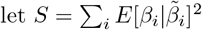. To define each bin, we first rank the posterior mean effect sizes based on their squared values 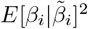. We define bin *b*_1_ as the smallest set of top SNPs with 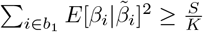 and iteratively define bin *b_k_* as the smallest set of additional top SNPs with 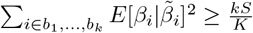. Let 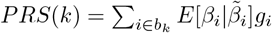. We define

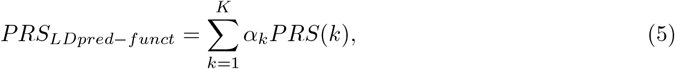

where the bin-specific weights *α_k_* are optimized using validation data via 10-fold cross-validation. For each held-out fold in turn, we split the data so we estimate the weights *α_k_* using the samples from the other nine folds (90% of the validation) and compute PRS on the held-out fold using these weights (10% of the validation). We then compute the average prediction *R*^2^ across the 10 held-out folds. To avoid overfitting when *K* is very close to *N*, we set the number of bins (*K*) to be between 1 and 100, such that it is proportional to 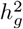 and the number of samples used to estimate the *K* weights in each fold is at least 100 times larger than *K*:

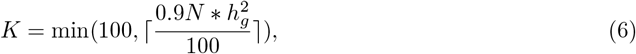

where *N* is the number of validation samples. For highly heritable traits 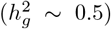, LDpred-funct reduces to the LDpred-funct-inf method if there are ~200 validation samples or fewer; for less heritable traits 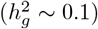, LDpred-funct reduces to the LDpred-funct-inf method if there are ~1,000 validation samples or fewer. In simulations, we set K to 40 (based on 7,585 validation samples; see below), approximately concordant with Equation 6. The value of 100 in the denominator of Equation 6 was coarsely optimized in simulations, but was not optimized using real trait data. We note that functional annotations are not used in the cross-validation step (although they do impact the posterior mean causal effect size provided as input to this step). Thus, it is likely that SNPs from a given functional annotation will fall into different bins (possibly all of the bins).

#### Standard errors

Standard errors for the prediction *R*^2^ of each method and the difference in prediction *R*^2^ between two methods were computed via block-jackknife using 200 genomic jackknife blocks^5^; this is more conservative than computing standard errors based on the number of validation samples, which does not account for variation across a finite number of SNPs. For each method, we first optimized any relevant tuning parameters using the entire genome and then analyzed each jackknife block using those tuning parameters.

### Simulations

We simulated quantitative phenotypes using real genotypes from the UK Biobank interim release (see below). We used up to 50,000 unrelated British-ancestry samples as training samples, and 7,585 samples of other European ancestries as validation samples (see below). We made these choices to minimize confounding due to shared population stratification or cryptic relatedness between training and validation samples (which, if present, could overstate the prediction accuracy that could be obtained in independent samples^36^), while preserving a large number of training samples. We restricted our simulations to 459,284 imputed SNPs on chromosome 1 (see below), fixed the number of causal SNPs at 2,000 or 5,000 (we also performed secondary simulations with 1,000 or 10,000 causal variants), and fixed the SNP-heritability 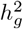 at 0.5. We sampled normalized causal effect sizes *β_i_* for causal SNPs from a normal distribution with variance equal to 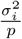, where *p* is the proportion of causal SNPs and 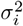 is the expected causal per-SNP heritability under the baseline-LD model^25^, fit using stratified LD score regression (S-LDSC)^5,25^ applied to height summary statistics computed from unrelated British-ancestry samples from the UK Biobank interim release (*N*=113,660). We computed per-allele effect sizes *b_i_* as 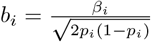, where *p_i_* is the minor allele frequency for SNP *i* estimated using the validation genotypes. We simulated phenotypes as 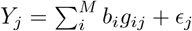, where 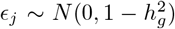. We set the training sample size to either 10,000, 20,000 or 50,000. The motivation to perform simulations using one chromosome is to be able to extrapolate performance at larger sample sizes^16^ according to the ratio *N/M*, where *N* is the training sample size. We compared each of the seven methods described above. For LDpred-funct-inf and LDpred-funct, for each simulated trait we used S-LDSC (applied to training data only) to estimate baseline-LD model parameters. For LDpred-funct, we report *R*^2^ as the average prediction *R*^2^ across the 10 held-out folds.

### Full UK Biobank data set

The full UK Biobank data set includes 459,327 European-ancestry samples and ~20 million imputed SNPs^30^ (after filtering as in ref. 27, excluding indels and structural variants). We selected 21 UK Biobank traits (14 quantitative traits and 7 binary traits) with phenotyping rate > 80% (> 80% of females for age at menarche, > 80% of males for balding), SNP-heritability 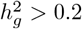 for quantitative traits, observed-scale SNP-heritability 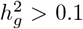 for binary traits, and low correlation between traits (as described in ref. 27). We restricted training samples to 409,728 British-ancestry samples^30^, including related individuals (avg *N*=373K phenotyped training samples; see Table S1 for quantitative traits and Table S2 for binary traits). We computed association statistics from training samples using BOLT-LMM v2.3^27^. We have made these association statistics publicly available (see Web Resources). We restricted validation samples to 24,436 samples of non-British European ancestry, after removing validation samples that were related (> 0.05) to training samples and/or other validation samples (avg *N*=22K phenotyped validation samples; see Table S1 and S2). As in our simulations, we made these choices to minimize confounding due to shared population stratification or cryptic relatedness between training and validation samples (which, if present, could overstate the prediction accuracy that could be obtained in independent samples^36^), while preserving a large number of training samples. We analyzed 6,334,603 genome-wide imputed SNPs, after removing SNPs with minor allele frequency < 1%, removing SNPs with imputation accuracy < 0.9, and removing A/T and C/G SNPs to eliminate potential strand ambiguity. We used 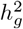 estimates from BOLT-LMM v2.3^27^ as input to LDpred, AnnoPred, LDpred-funct-inf and LDpred-funct.

### UK Biobank interim release

The UK Biobank interim release includes 145,416 European-ancestry samples^37^. We used the UK Biobank interim release both in simulations using real genotypes, and in a subset of analyses of height phenotypes (to investigate how prediction accuracy varies with training sample size).

In our analyses of height phenotypes, we restricted training samples to 113,660 unrelated (≤ 0.05) British-ancestry samples for which height phenotypes were available. We computed association statistics by adjusting for 10 PCs^38^, estimated using FastPCA^39^ (see Web Resources). For consistency, we used the same set of 24,351 validation samples of non-British European ancestry with height phenotypes as defined above. We analyzed 5,957,957 genome-wide SNPs, after removing SNPs with minor allele frequency < 1%, removing SNPs with imputation accuracy < 0.9, removing SNPs that were not present in the 23andMe height data set (see below), and removing A/T and C/G SNPs to eliminate potential strand ambiguity.

In our simulations, we restricted training samples to up to 50,000 of the 113,660 unrelated British-ancestry samples, and restricted validation samples to 8,441 samples of non-British European ancestry, after removing validation samples that were related (> 0.05) to training samples and/or other validation samples. We restricted the 5,957,957 genome-wide SNPs (see above) to chromosome 1, yielding 459,284 SNPs after QC.

### 23andMe height summary statistics

The 23andMe data set consists of summary statistics computed from 698,430 European-ancestry samples (23andMe customers who consented to participate in research) at 9,898,287 imputed SNPs, after removing SNPs with minor allele frequency < 1% and that passed QC filters (which include filters on imputation quality, avg.rsq< 0.5 or min.rsq< 0.3 in any imputation batch, and imputation batch effects). Analyses were restricted to the set of individuals with > 97% European ancestry, as determined via an analysis of local ancestry^40^. Summary association statistics were computed using linear regression adjusting for age, gender, genotyping platform, and the top five principal components to account for residual population structure. The summary association statistics will be made available to qualified researchers (see Web Resources).

We analyzed 5,808,258 genome-wide SNPs, after removing SNPs with minor allele frequency < 1%, removing SNPs with imputation accuracy < 0.9, removing SNPs that were not present in the full UK Biobank data set (see above), and removing A/T and C/G SNPs to eliminate potential strand ambiguity.

### Meta-analysis of full UK Biobank and 23andMe height data sets

We meta-analyzed height summary statistics from the full UK Biobank and 23andMe data sets. We define

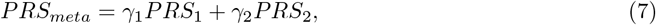

where *PRS_i_* is the PRS obtained using training data from cohort *i*. The PRS can be obtained using P+T, P+T-funct-LASSO, LDpred-inf or LDpred-funct. The meta-analysis weights γ*_i_* can either be specified via fixed-effect meta-analysis (e.g. 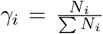) or optimized using validation data^26^. We use the latter approach, which can improve prediction accuracy (e.g. if the cohorts differ in their heritability as well as their sample size). In our primary analyses, we fit the weights γ*_i_* in-sample and report prediction accuracy using adjusted *R*^2^ to account for in-sample fitting^26^. We also report results using 10-fold cross-validation: for each held-out fold in turn, we estimate the weights γ*_i_* using the other nine folds and compute PRS on the held-out fold using these weights. We then compute the average prediction *R*^2^ across the 10 held-out folds.

When using LDpred-funct as the prediction method, we perform the meta-analysis as follows. First, we use LDpred-funct-inf to fit meta-analysis weights γ*_i_*. Then, we use γ*_i_* to compute (metaanalysis) weighted posterior mean causal effect sizes (PMCES) via *PMCES* = *α*_1_*PMCES*_1_ + *α*_2_*PMCES*_2_, which are binned into *k* bins. Then, we estimate bin-specific weights *α_k_* (used to compute (meta-analysis + bin-specific) weighted posterior mean causal effect sizes 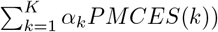 using validation data via 10-fold cross validation.

### Baseline-LD model annotations

The baseline-LD model (v1.1) contains a broad set of 75 functional annotations (including coding, conserved, regulatory and LD-related annotations), whose enrichments are jointly estimated using stratified LD score regression^5,25^. For each trait, we used the *τ_c_* values estimated for that trait to compute 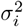, the expected per-SNP heritability of SNP *i* under the baseline-LD model, as

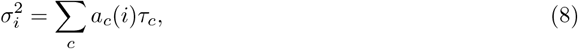

where *a_c_*(*i*) is the value of annotation *c* at SNP *i*.

Joint effect sizes *τ_c_* for each annotation *c* are estimated via

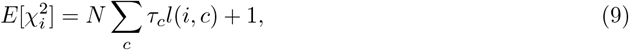

where *l*(*i, c*) is the LD score of SNP *i* with respect to annotation *a_c_* and 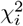 is the chi-square statistic for SNP *i*. We note that *τ_c_* quantifies effects that are unique to annotation *c*. In all analyses of real phenotypes, *τ_c_* and 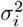 were estimated using training samples only.

In our primary analyses, we used 489 unrelated European samples from phase 3 of the 1000 Genomes Project^33^ as the reference data set to compute LD scores, as in ref. 25.

To verify that our 1000 Genomes reference data set produces reliable LD estimates, we repeated our LDpred-funct analyses using S-LDSC with 3,567 unrelated individuals from UK10K^41^ as the reference data set (as in ref. 42), ensuring a closer ancestry match with British-ancestry UK Biobank samples. We also repeated our LDpred-funct analyses using S-LDSC with the baseline-LD+LDAK model (instead of the baseline-LD model), with UK10K as the reference data set. The baseline-LD+LDAK model (introduced in ref. 42) consists of the baseline-LD model plus one additional continuous annotation constructed using LDAK weights^43^, which has values (*p_j_*(1 − *p_j_*))^1+*α*^ *w_j_*, where *α* = −0.25, *p_j_* is the allele frequency of SNP *j*, and *w_j_* is the LDAK weight of SNP *j* computed using UK10K data.

## Results

### Simulations

We performed simulations using real genotypes from the UK Biobank interim release and simulated phenotypes (see Methods). We simulated quantitative phenotypes with SNP-heritability 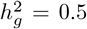, using 476,613 imputed SNPs from chromosome 1. We selected either 2,000 or 5,000 variants to be causal; we refer to these as “sparse” and “polygenic” architectures, respectively. We sampled normalized causal effect sizes from normal distributions with variances based on expected causal per-SNP heritabilities under the baseline-LD model^25^, fit using stratified LD score regression (S-LDSC)^5,25^ applied to height summary statistics from British-ancestry samples from the UK Biobank interim release. We randomly selected 10,000, 20,000 or 50,000 unrelated British-ancestry samples as training samples, and we used 7,585 unrelated samples of non-British European ancestry as validation samples. By restricting simulations to chromosome 1 (≈ 1/10 of SNPs), we can extrapolate results to larger sample sizes (≈ 10x larger; see Application to 21 UK Biobank traits), analogous to previous work^16^.

We compared prediction accuracies (*R*^2^) for seven main methods: P+T^18,19^, LDpred^16^, SBayesR^9^, P+T-funct-LASSO^23^, AnnoPred^24^, LDpred-funct-inf and LDpred-funct (see Methods). Results are reported in Figure 1 (main simulations) and Figure S1 (additional values of number of causal variants); numerical results are reported in Table S3 and Table S4. Among methods that do not use functional information, the prediction accuracy of LDpred was higher than P+T (particularly for the polygenic architecture), consistent with previous work^8,16^ (see Table S5 and Table S6 for optimal tuning parameters; surprisingly, at *N*=50K training samples, LDpred is optimized by assuming that 100% of SNPs are causal). SBayesR attained a substantial improvement vs. LDpred at *N*=10K training samples (+19% relative improvement for sparse architecture and +8.6% relative improvement for polygenic architecture) but attained prediction *R*^2^ close to 0 at larger sample sizes (*N*=20K and *N*=50K), perhaps because the algorithm failed to converge (Table S3; results not included in Figure 1).

**Figure 1:**
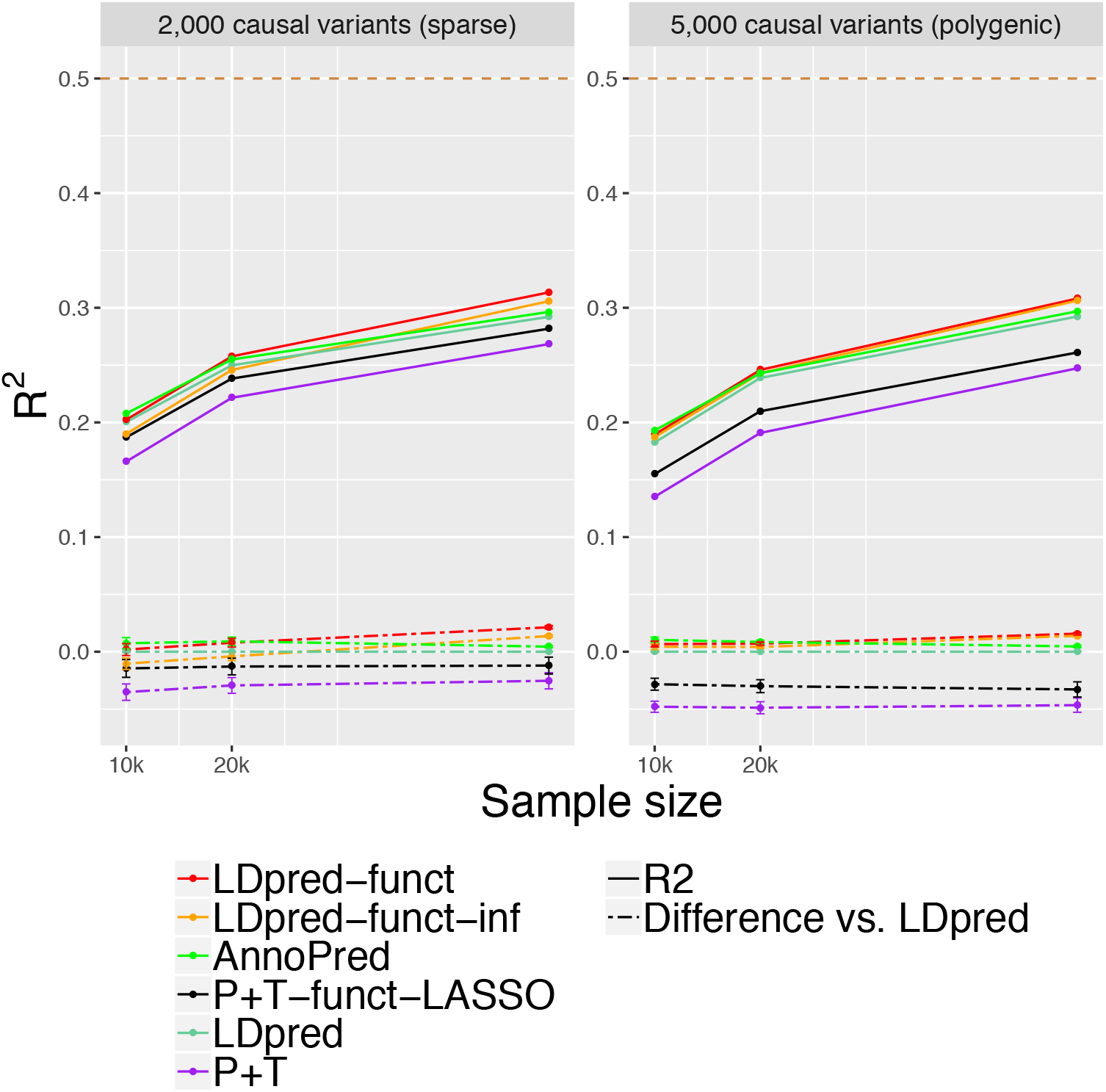
Accuracy of 6 polygenic prediction methods in simulations using UK Biobank genotypes. We report results for P+T, LDpred, P+T-funct-LASSO, AnnoPred, LDpred-funct-inf and LDpred-funct in chromosome 1 simulations with 2,000 causal variants (sparse architecture) and 5,000 causal variants (polygenic architecture). Results are averaged across 100 simulations. Top dashed line denotes simulated SNP-heritability of 0.5. Bottom dashed lines denote differences vs. LDpred; error bars represent 95% confidence intervals. Results for other values of the number of causal variants are reported in Figure S1, and numerical results are reported in Table S3 and Table S4.

Incorporating functional information via LDpred-funct-inf (a method that does not model sparsity) produced improvements that varied with sample size (+4.7% relative improvement for sparse architecture and +4.8% relative improvement for polygenic architecture at *N*=50K training samples, compared to LDpred; smaller improvements at smaller sample sizes). These results are consistent with the fact that LDpred is known to be sensitive to model assumptions at large sample sizes^16^. Accounting for sparsity using LDpred-funct further improved prediction accuracy, particularly for the sparse architecture (+7.3% relative improvement for sparse architecture and +5.4% relative improvement for polygenic architecture at *N*=50K training samples, compared to LDpred; smaller improvements at smaller sample sizes). LDpred-funct attained substantially higher prediction accuracy than P+T-funct-LASSO in most settings (+11% relative improvement for sparse architecture and +18% relative improvement for polygenic architecture at *N*=50K training samples; smaller improvements at smaller sample sizes). LDpred-funct also attained higher prediction accuracy than AnnoPred at large sample sizes (+5.7% relative improvement for sparse architecture and +3.7% relative improvement for polygenic architecture at *N*=50K training samples; smaller differences at smaller sample sizes) (see Table S7 for optimal tuning parameters; surprisingly, at *N*=50K training samples, AnnoPred is optimized by assuming that 100% of SNPs are causal, analogous to LDpred). The difference in prediction accuracy between LDpred and each other method, as well as the difference in prediction accuracy between LDpred-funct and each other method, was statistically significant in most cases (see Table S4 e.g. vs. AnnoPred: *P* < 10^−125^ for sparse architecture and *P* < 10^−75^ for polygenic architecture at *N*=50K training samples). Simulations with 1,000 or 10,000 causal variants generally recapitulated these findings, although SBayesR, P+T-funct-LASSO and AnnoPred performed better than LDpred-funct for the extremely sparse architecture at N=10K (Table S3).

The average running time for all 7 methods is reported in Table S8. We separately report the time to estimate posterior mean causal effect sizes, and the time to compute LD matrices (not applicable for LDpred-funct-inf and LDpred-funct) (we do not include the time to compute polygenic risk scores, which is small in comparison and depends on the number of validation samples). For the two methods with highest prediction *R*^2^ in analyses of real UK Biobank traits (LDpred-funct and AnnoPred; see below), the average running time was 71 minutes for LDpred-funct vs. 5,249 minutes for AnnoPred, not including the time to compute LD matrices.

We performed four secondary analyses. First, we assessed the calibration of each method by checking whether a regression of true vs. predicted phenotype yielded a slope of 1. We determined that LDpred-funct was well-calibrated (regression slope 0.98-0.99), LDpred and AnnoPred were fairly well-calibrated (regression slope 0.85-1.00), and other methods were not well-calibrated (Table S9). Second, we assessed the sensitivity of LDpred-funct to the choice of *K*=40 posterior mean causal effect size bins to regularize effect sizes in our main simulations. We determined that results were not sensitive to this parameter (Table S10); slightly higher values of *K* performed slightly better, but we did not finely optimize this parameter. Third, we evaluated a “cheating” version of LDpred-funct that utilized the true baseline-LD model parameters used to simulate the data, instead of estimating these parameters from the data (LDpred-funct-cheat). LDpred-funct-cheat performed only slightly better than LDpred-funct, indicating that LDpred-funct is not sensitive to imperfect estimation of functional enrichment parameters (see Table S11). Fourth, we simulated traits with lower SNP-heritability 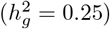 (see Table S12). We determined that the improvements attained by LDpred-funct were smaller in these simulations (e.g. +6.9% relative improvement vs. AnnoPred and −1.0% relative improvement vs. LDpred for sparse architecture, +3.4% improvement vs. AnnoPred and +0.6% relative improvement vs. LDpred for polygenic architecture at *N*=50K training samples; smaller improvements at smaller sample sizes).

### Application to 21 UK Biobank traits

We applied P+T, LDpred, SBayesR, P+T-funct-LASSO, AnnoPred, LDpred-funct-inf and LDpred-funct to 21 UK Biobank traits (14 quantitative traits and 7 binary traits; Table S1 and Table S2). We analyzed training samples of British ancestry (avg *N*=373K) and validation samples of non-British European ancestry (avg *N*=22K). We included 6,334,603 imputed SNPs in our analyses (see Methods). We computed summary statistics and 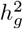 estimates from training samples using BOLT-LMM v2.3^27^ (see Table S13). We estimated trait-specific functional enrichment parameters for the baseline-LD model^25^ by running S-LDSC^5,25^ on these summary statistics. Results for quantitative traits are reported in Figure 2 and Table S14, and results for binary traits are reported in Figure 3 and Table S15. Differences between each main prediction method and either LDpred or LDpred-funct (and block-jackknife standard errors on these differences) are reported in Table S16, and averages across all 21 traits for main and secondary prediction methods are reported in Table S17.

**Figure 2:**
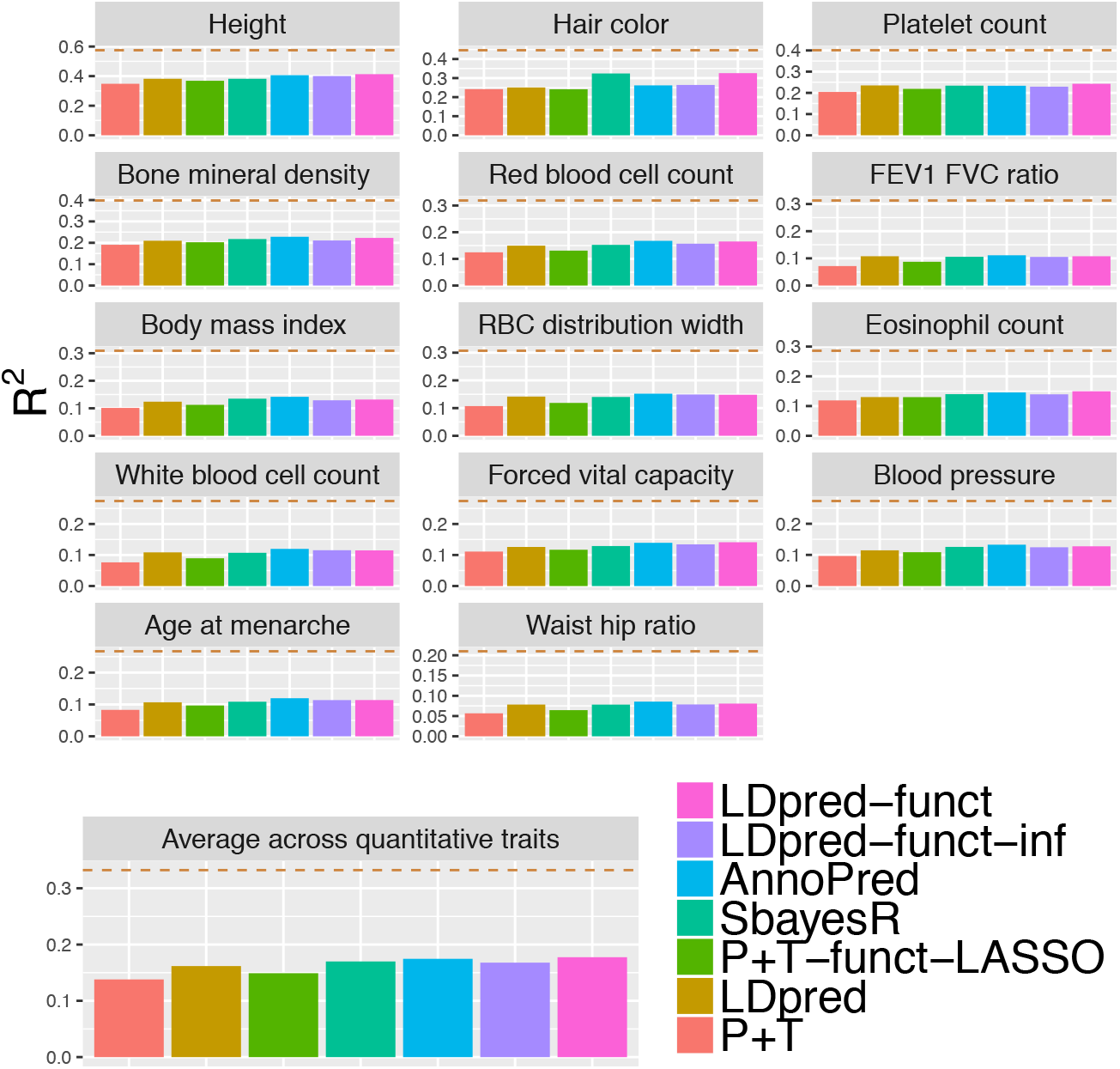
Accuracy of 7 polygenic prediction methods across 14 UK Biobank quantitative traits. We report results for P+T, LDpred, SBayesR, P+T-funct-LASSO, AnnoPred, LDpred-funct-inf and LDpred-funct. Dashed lines denote estimates of SNP-heritability. Numerical results are reported in Table S14.

**Figure 3:**
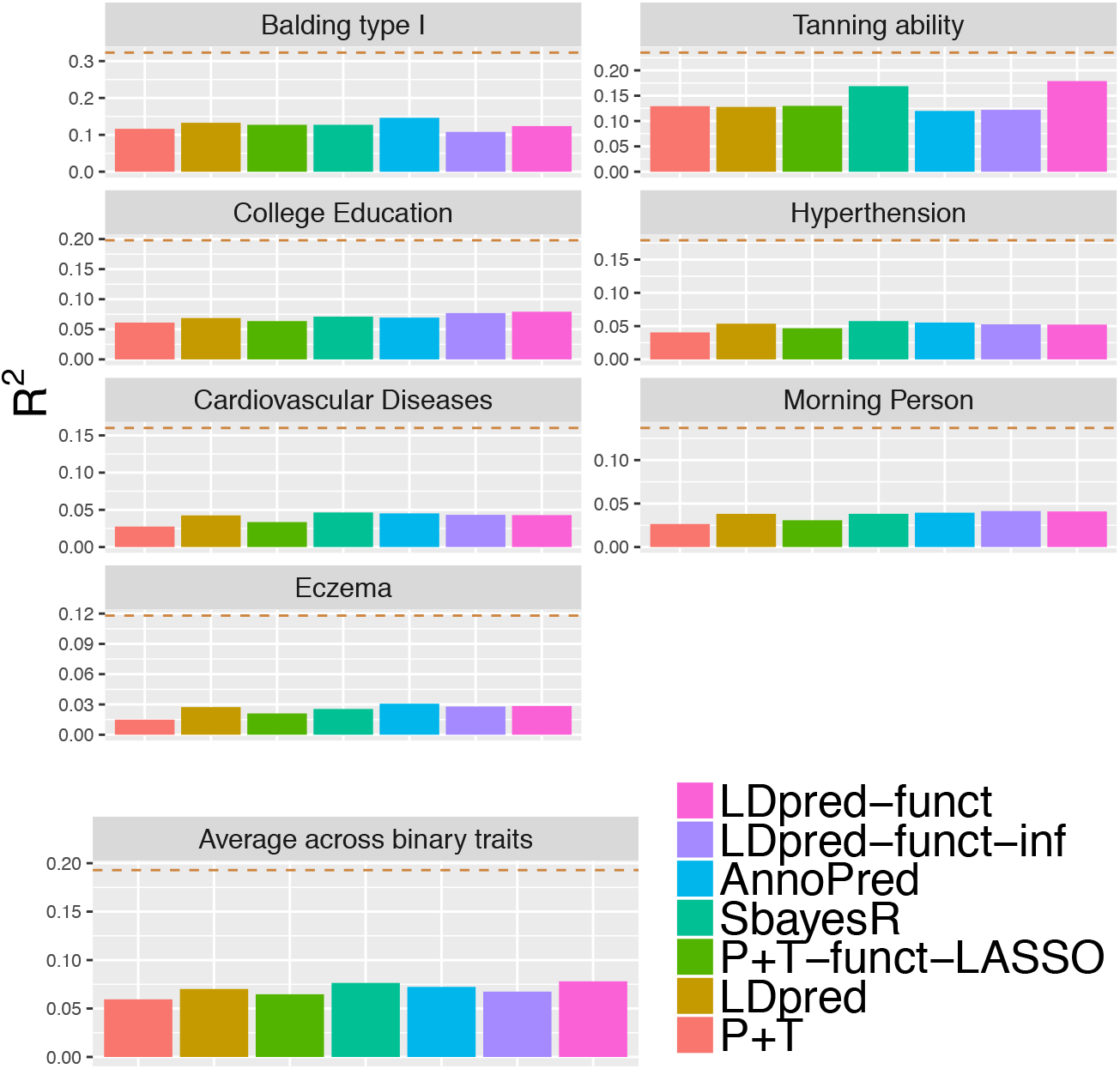
Accuracy of 7 polygenic prediction methods across 7 UK Biobank binary traits. We report results for P+T, LDpred, SBayesR, P+T-funct-LASSO, AnnoPred, LDpred-funct-inf and LDpred-funct. Dashed lines denote estimates of SNP-heritability. Numerical results are reported in Table S15.

Among methods that do not use functional information, LDpred outperformed P+T (+18% relative improvement in average prediction *R*^2^), consistent with simulations under a polygenic architecture (see Table S18 and Table S19 for optimal tuning parameters) and with previous work^8,16^. LDpred also outperformed LDpred-inf, a method that does not model sparsity (see Table S17). The exclusion of long-range LD regions (see Methods) was critical to LDpred performance, as running LDpred without excluding long-range LD regions (as implemented in a previous version of this paper^44^) performed much worse (see Table S17). SBayesR outperformed LDpred (+5.3% relative improvement in average prediction *R*^2^), with no convergence issues in the full UK Biobank analysis (but see below for 113K interim UK Biobank analysis); we note that expanding the set of SNPs analyzed worsened the performance of SBayesR (see below).

Incorporating functional information via LDpred-funct-inf (a method that does not model sparsity) performed only slightly better than LDpred (+0.9% improvement in average prediction *R*^2^), but greatly outperformed LDpred-inf (+19% relative improvement, *P* < 10^−20^ for difference using two-sided z-test based on block-jackknife standard error in Table S20). Accounting for sparsity using LDpred-funct substantially improved prediction accuracy (+10%, +4.6%, +7.4% relative improvements in average prediction *R*^2^ vs. LDpred, SBayesR, LDpred-funct-inf; *P* < 2 * 10^−4^, *P* = 0.04, *P* < 2 * 10^−4^ for differences using two-sided z-test based on block-jackknife standard error in Table S16; average prediction *R*^2^=0.144; highest *R*^2^=0.413 for height), consistent with simulations. The relative improvement in avg prediction *R*^2^ for LDpred-funct vs. LDpred was +9.7% for quantitative traits (higher prediction *R*^2^ for 14/14 traits), and +11% for binary traits (higher prediction *R*^2^ for 5/7 traits). We observed a positive but non-significant correlation across traits between /E and relative improvement (Figure S2), perhaps due to the limited number of data points and/or contribution of other factors (e.g. polygenicity). LDpred-funct also performed substantially better than P+T-funct-LASSO (+20% relative improvement in avg. prediction *R*^2^), consistent with simulations under a polygenic architecture. AnnoPred performed slightly but non-significantly worse than LDpred-funct (−2.7% relative change in average prediction *R*^2^ for AnnoPred vs. LDpred-funct, *P* = 0.35 for difference using two-sided z-test based on block-jackknife standard error in Table S16; see Table S21 for optimal tuning parameters).

In the above experiments, LDpred-funct analyzed 373K training samples and 22K validation samples and used 90% of the validation samples to estimate regularization weights. It is possible that incorporating data from an additional 20K samples could confer an unfair advantage for LDpred-funct compared to other methods. To assess this, we performed two new experiments. First, we repeated the LDpred-funct analyses using smaller validation sample sizes (as low as 1K). We determined that results were little changed (Table S22). Second, we repeated the LDpred-funct analyses using all 22K validation samples but using only 1K samples to estimate validation weights. Again, we determined that results were little changed (Table S23). As the use of 1K samples to estimate validation weights is a trivial number of additional samples compared to 373K training samples, we conclude from these experiments that LDpred-funct does not owe its advantage to incorporating data from a substantial number of additional samples.

We performed 13 secondary analyses. First, we assessed the calibration of each method by checking whether a regression of true vs. predicted phenotype yielded a slope of 1. As in our simulations, we determined that LDpred-funct was well-calibrated (average regression slope: 0.98), LDpred and AnnoPred were fairly well-calibrated (average regression slope: 0.89 and 0.83, respectively), and other methods were not well-calibrated (Table S24). Second, we assessed the sensitivity of LDpred-funct to the average value of *K* = 58 posterior mean causal effect size bins to regularize effect sizes in these analyses (see Equation 6 and Table S13). We determined that results were not sensitive to the number of bins (Table S25). Third, we determined that functional enrichment information is far less useful when restricting to genotyped variants (e.g. −6.9% relative change in avg prediction *R*^2^ for LDpred-funct vs. LDpred when both methods are restricted to typed variants; Table S17), likely because tagging variants may not belong to enriched functional annotations. Fourth, we repeated the SBayesR analysis using the 2.9M SNP set instead of the 1.1M SNP set (see Methods), but determined that this substantially worsened the performance of SBayesR (Table S17). Fifth, we evaluated a modification of P+T-funct-LASSO in which different weights were allowed for the two predictors (P+T-funct-LASSO-weighted; see Methods), but results were little changed (+1.1% relative improvement in avg prediction *R*^2^ vs. P+T-funct-LASSO; Table S17). Sixth, we obtained similar results for P+T-funct-LASSO when defining the “high-prior” (HP) SNP set using the top 5% of SNPs with the highest per-SNP heritability, instead of the top 10% (see Table S17). Seventh, we determined that incorporating baseline-LD model functional enrichments that were meta-analyzed across traits (31 traits from ref. 25), instead of the trait-specific functional enrichments used in our primary analyses, slightly reduced the prediction accuracy of LDpred-funct-inf (Table S17). Eigth, to assess whether the improvement of LDpred-funct is specific to the 75 functional annotations of the baseline-LD model, we implemented an analogous method that uses 75 random annotations (LDpred-funct (random)). We determined that LDpred-funct attained a 13% relative improvement in average prediction *R*^2^ vs. LDpred-funct (random), which performed similarly to LDpred (3.1% decrease in average prediction *R*^2^ vs. LDpred) (Table S17). This implies that the improvement of LDpred-funct is specific to the 75 functional annotations of the baseline-LD model. We further note that a method that does not use functional priors but applies the regularization step of LDpred-funct on top of LDpred-inf (LDpred-inf + sparsity) performed similarly to LDpred-funct (random) (Table S17).Ninth, we determined that using our previous baseline model^5^, instead of the baseline-LD model^25^, slightly reduced the prediction accuracy of LDpred-funct-inf and LDpred-funct (Table S17). Tenth, to assess how much of the improvement of LDpred-funct derives from the removal of uninformative SNPs, we implemented an analogous method that uses functional annotations to restrict to the same set of SNPs with expected per-SNP heritability 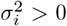 (2,981,166-4,306,498 SNPs depending on the trait; see Methods) but then imposes a constant prior on causal effect sizes (LDpred-funct (constant prior)). We determined that LDpred-funct attained a 4.3% relative improvement in average prediction *R*^2^ vs. LDpred-funct (constant prior), which attained a 5.5% relative improvement in average prediction *R*^2^ vs. LDpred (Table S17). Thus, some but not all of the improvement of LDpred-funct derives from the removal of (relatively) uninformative SNPs. Eleventh, we determined that inferring functional enrichments using only the SNPs that passed QC filters and were used for prediction had no impact on the prediction accuracy of LDpred-funct-inf (Table S17). Twelveth, we determined that using UK10K (instead of 1000 Genomes) as the LD reference panel had virtually no impact on prediction accuracy (Table S17). Thirteenth, we determined that using UK10K (instead of 1000 Genomes) as the LD reference panel had virtually no impact on prediction accuracy (Table S17).

### Application to height in meta-analysis of UK Biobank and 23andMe cohorts

We applied P+T, LDpred-inf, SBayesR, P+T-funct-LASSO, AnnoPred, LDpred-funct-inf and LDpred-funct to predict height in a meta-analysis of UK Biobank and 23andMe cohorts (see Methods). Training sample sizes were equal to 408,092 for UK Biobank and 698,430 for 23andMe, for a total of 1,106,522 training samples. For comparison purposes, we also computed predictions using the UK Biobank and 23andMe training data sets individually, as well as a training data set consisting of 113,660 British-ancestry samples from the UK Biobank interim release. (The analysis using the 408,092 UK Biobank training samples was nearly identical to the analysis of Figure 2, except that we used a different set of 5,957,935 SNPs, for consistency throughout this set of comparisons; see Methods.) We used 24,351 UK Biobank samples of non-British European ancestry as validation samples in all analyses.

Results are reported in Figure 4 and Table S26. The relative improvements attained by LDpred-funct-inf and LDpred-funct were broadly similar across all four training data sets (also see Figure 2), implying that these improvements are not specific to the UK Biobank data set. Interestingly, compared to the full UK Biobank training data set (*R*^2^=0.415 for LDpred-funct; slightly different from *R*^2^ = 0.413 in Figure 2 due to slightly different SNP set), prediction accuracies were only slightly higher for the meta-analysis training data set (*R*^2^=0.431 for LDpred-funct), and were lower for the 23andMe training data set (*R*^2^=0.344 for LDpred-funct), consistent with the ≈ 30% higher heritability in UK Biobank as compared to 23andMe and other large cohorts^25,27,28^; the higher heritability in UK Biobank could potentially be explained by lower environmental heterogeneity. We note that in the meta-analysis, we optimized the meta-analysis weights using validation data (similar to ref. 26), instead of performing a fixed-effect meta-analysis. This approach accounts for differences in heritability as well as sample size, and attained a +3.3% relative improvement in prediction *R*^2^ compared to fixed-effects meta-analysis (see Table S26). We note that SBayesR performed similarly to LDpred in height analyses with ≥ 408K training samples (−10% to +0.2% change in average prediction *R*^2^) but attained prediction *R*^2^ close to 0 in the height analysis with 113K training samples, perhaps because the algorithm failed to converge (Table S26; results not included in Figure 4).

**Figure 4:**
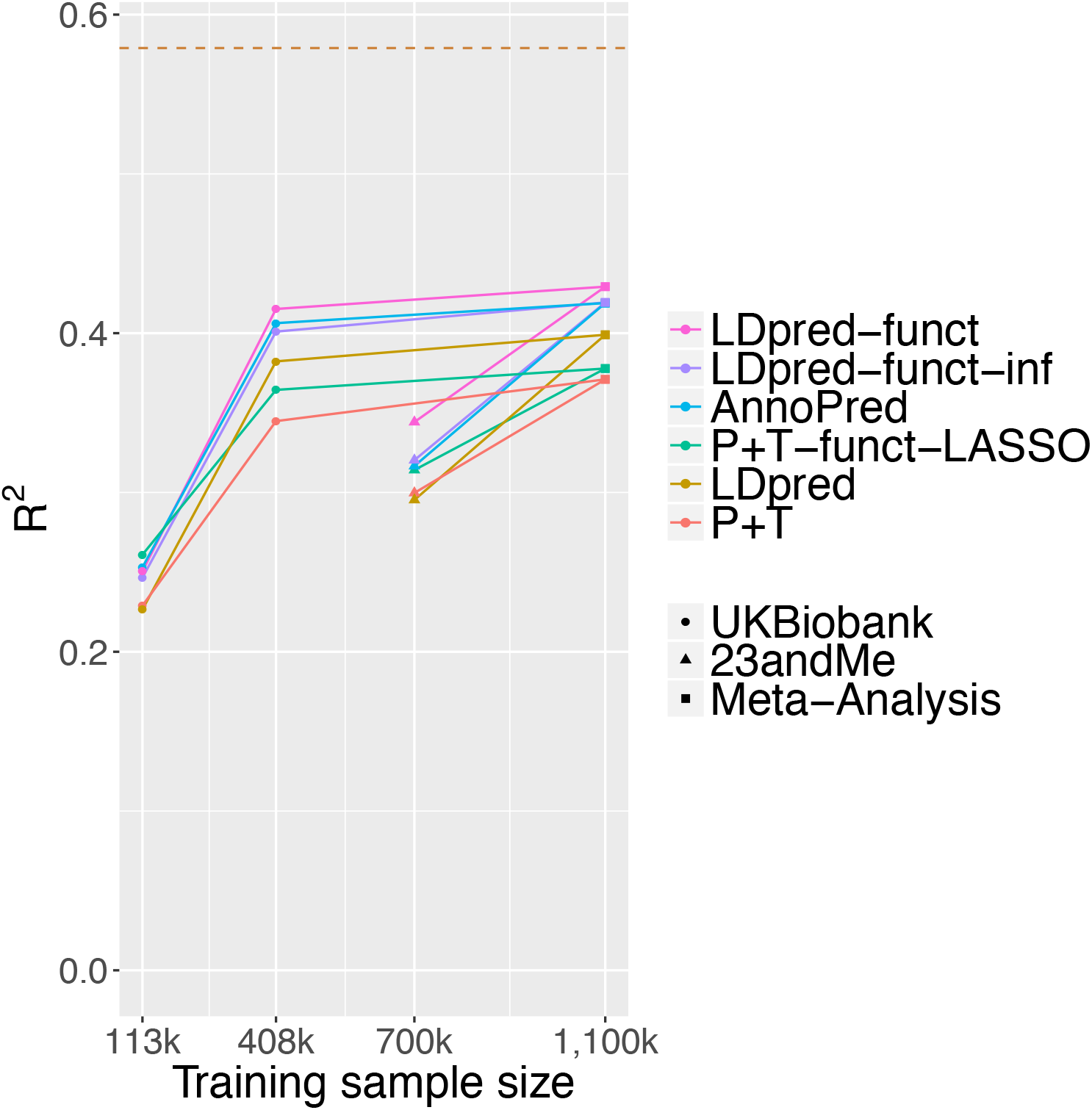
Accuracy of 6 prediction methods in height meta-analysis of UK Biobank and 23andMe cohorts. We report results for P+T, LDpred, P+T-funct-LASSO, AnnoPred, LDpred-funct-inf and LDpred-funct, for each of 4 training data sets: UK Biobank interim release (113,660 training samples), UK Biobank (408,092 training samples), 23andMe (698,430 training samples) and meta-analysis of UK Biobank and 23andMe (1,107,430 training samples). Nested training data sets are connected by solid lines (e.g. UK Biobank (408k) and 23andMe are both connected to Meta-Analysis, but not to each other). Dashed line denotes estimate of SNP-heritability in UK Biobank. Numerical results are reported in Table S26.

## Discussion

We have shown that leveraging trait-specific functional enrichments inferred by S-LDSC with the baseline-LD model^25^ substantially improves polygenic prediction accuracy. Across 21 UK Biobank traits, we attained substantial improvements in average prediction *R*^2^ using a method that leverages functional enrichment and performs an additional regularization step to account for sparsity (LDpred-funct). LDpred-funct attained +10% (*P* < 2 * 10^−4^) and +4.6% (*P* = 0.04) relative improvements compared to LDpred^16^ and SBayesR^9^, two state-of-the-art methods that do not model functional enrichment. Thus incorporating functional annotations improves polygenic prediction accuracy. We note that our main analyses used baseline-LD model v1.1, but using the updated baseline-LD model v2.1 yields slightly higher prediction *R*^2^ for LDpred-funct-inf and LDpred-funct (Table S17).

Two previous studies have highlighted the potential advantages of leveraging functional enrichment to improve prediction accuracy^23,24^. We included both of these methods in all of our analyses. First, ref. 23 introduced a method (which we call P+T-funct-LASSO) that corrects marginal effect sizes for winner’s curse using LASSO and incorporates functional data to define high-prior and low-prior SNP sets. LDpred-funct attained a +19% average relative improvement vs. P+T-funct-LASSO across 21 UK Biobank traits. Second, ref. 24 introduced AnnoPred, which uses a Bayesian framework to incorporate functional annotations. AnnoPred models sparsity differently than LDpred-funct, as it uses a point-normal prior to estimate posterior mean effect sizes via Markov Chain Monte Carlo (MCMC), whereas LDpred-funct performs a regularization step to account for sparsity. We note that ref. 24 considered only genotyped variants and binary annotations. As noted above, functional enrichment information is far less useful when restricting to genotyped variants (Table S17), likely because tagging variants may not belong to enriched functional annotations; thus, the utility of AnnoPred in more general settings is currently unknown. Here, we determined that AnnoPred performed slightly but non-significantly worse than LDpred-funct(−2.3% relative change in average prediction *R*^2^; *P* = 0.35 for difference) across 21 UK Biobank traits, consistent with slightly worse results for AnnoPred in simulations at large sample sizes. We emphasize that our current work represents, to our knowledge, the first effort to combine binary and continuous-valued functional annotations to improve polygenic risk prediction using imputed variants.

Our work has several limitations. First, LDpred-funct analyzes summary statistic training data (which are publicly available for a broad set of diseases and traits^45^), but methods that use raw genotypes/phenotypes as training data have the potential to attain higher accuracy^27^; incorporating functional enrichment information into prediction methods that use raw genotypes/phenotypes as training data remains a direction for future research. Second, the regularization step employed by LDpred-funct to account for sparsity relies on heuristic cross-validation instead of inferring posterior mean causal effect sizes under a prior sparse functional model; we made this choice because the appropriate choice of sparse functional model is unclear, and because inference of posterior means via MCMC may be subject to convergence issues. As a consequence, the improvement of LDpred-funct over LDpred-funct-inf may be contingent on the number of validation samples available for crossvalidation; in particular, for very small validation samples, the number of cross-validation bins is equal to 1 (Equation 6) and LDpred-funct is identical to LDpred-funct-inf. However, we determined that results of LDpred-funct were little changed when restricting to smaller validation sample sizes (as low as 1,000; see Table S22) or using all 22K validation samples but using only 1K samples to estimate validation weights (Table S23); this implies that LDpred-funct does not owe its advantage to incorporating data from a substantial number of additional samples. Third, we have considered only single-trait analyses, but leveraging genetic correlations among traits has considerable potential to improve prediction accuracy^46,47^. Fourth, we have not considered how to leverage functional enrichment for polygenic prediction in related individuals^48^. Fifth, we have not thoroughly investigated the application LDpred-funct to polygenic prediction in diverse populations^26,49–51^ (for which very similar functional enrichments have been reported^52,53^), as our simulations focused exclusively on prediction in Europeans. However, we evaluated the performance of LDpred-funct in predicting 21 UK Biobank traits in diverse populations using European training data (as in recent studies^49,50^). The results were promising, particularly in Africans (+23% vs. LDpred (P < 10^−5^), +18% vs. SBayesR (P = 0.001); see Table S27), for which distinguishing causal vs. non-causal variants is particularly important due to differences in LD vs. Europeans^54^. A more thorough investigation, e.g. incorporating non-European training data^26^, is an important direction for future research. Sixth, we have not performed a comprehensive assessment of how much different functional annotation models contribute to improvements in prediction accuracy, which remains as an important future direction, particularly as functional annotation models will improve as increasingly rich functional data is generated. Specifically, the improvements in prediction accuracy that we reported are a function of the baseline-LD model^25^, but there are many possible ways to improve this model, e.g. by incorporating tissue-specific enrichments^1–6,55–58^, modeling MAF-dependent architectures^59–61^, and/or employing alternative approaches to modeling LD-dependent effects^43^; we anticipate that future improvements to the baseline-LD model will yield even larger improvements in prediction accuracy. As an initial step to explore alternative approaches to modeling LD-dependent effects, we repeated our analyses using the baseline-LD+LDAK model (introduced in ref. 42), which consists of the baseline-LD model plus one additional continuous annotation constructed using LDAK weights^43^. (Recent work has shown that incorporating LDAK weights increases polygenic prediction accuracy in analyses that do not include the baseline-LD model^62^.) We determined that results were virtually unchanged (avg prediction *R*^2^=0.1350 for baseline-LD+LDAK vs. 0.1354 for baseline-LD using LDpred-funct-inf with UK10K SNPs; see Table S17 and Table S28). Despite these limitations and open directions for future research, our work demonstrates that leveraging functional enrichment using the baseline-LD model substantially improves polygenic prediction accuracy.

## Acknowledgements

We thank the research participants and employees of 23andMe for making this work possible. We are grateful to S. Sunyaev, S. Chun, L. O’Connor, O. Weissbrod and H. Finucane for helpful discussions. This research was conducted using the UK Biobank Resource under Application #16549 and was funded by NIH grants R01 GM105857, R01 MH101244 and U01 HG009379.

Collaborators for the 23andMe research team are: Michelle Agee, Babak Alipanahi, Robert K. Bell, Katarzyna Bryc, Sarah L. Elson, Pierre Fontanillas, David A. Hinds, Jennifer C. McCreight, Karen E. Huber, Aaron Kleinman, Nadia K. Litterman, Matthew H. McIntyre, Joanna L. Mountain, Elizabeth S. Noblin, Carrie A.M. Northover, Steven J. Pitts, J. Fah Sathirapongsasuti, Olga V. Sazonova, Janie F. Shelton, Suyash Shringarpure, Chao Tian, Joyce Y. Tung, Vladimir Vacic, and Catherine H. Wilson.

## Author contributions

C.M.L. and A.L.P. designed experiments. C.M.L. performed experiments. C.M.L., S.G., P.R.L., S.S.K., N.F. and A.A. analyzed data. C.M.L. and A.L.P. wrote the manscript with assistance from S.G., P.R.L. S.S.K., N.F. and A.L.P.

## Web Resources

Software implementing the LDpred-funct-inf and LDpred-funct: https://www.hsph.harvard.edu/alkes-price/software

LDscore regression software: https://github.com/bulik/ldsc

UK Biobank Resource: http://www.ukbiobank.ac.uk/

BOLT-LMM v2.3 software http://data.broadinstitute.org/alkesgroup/BOLT-LMM/

BOLT-LMM v2.3 association statistics: https://data.broadinstitute.org/alkesgroup/UKBB/UKBB_409K/

23andMe height association statistics: The full summary statistics for the 23andMe height GWAS will be made available through 23andMe to qualified researchers under an agreement with 23andMe that protects the privacy of the 23andMe participants. Please visit https://research.23andme.com/collaborate/#publication for more information and to apply to access the data.

AnnoPred: https://github.com/yiminghu/AnnoPred

SBayesR software: http://cnsgenomics.com/software/gctb/. SBayesR shrunk and sparse LD matrices can be downloaded from Zenodo public repository, for both 1.09 million HapMap3 (10.5281/zenodo.3350914) and 2.8 million pruned variants (10.5281/zenodo.3375373).

