## Supplementary Material for "LDpred-funct: incorporating functional priors improves polygenic prediction accuracy in UK Biobank and 23andMe data sets"

### Supplementary Figures

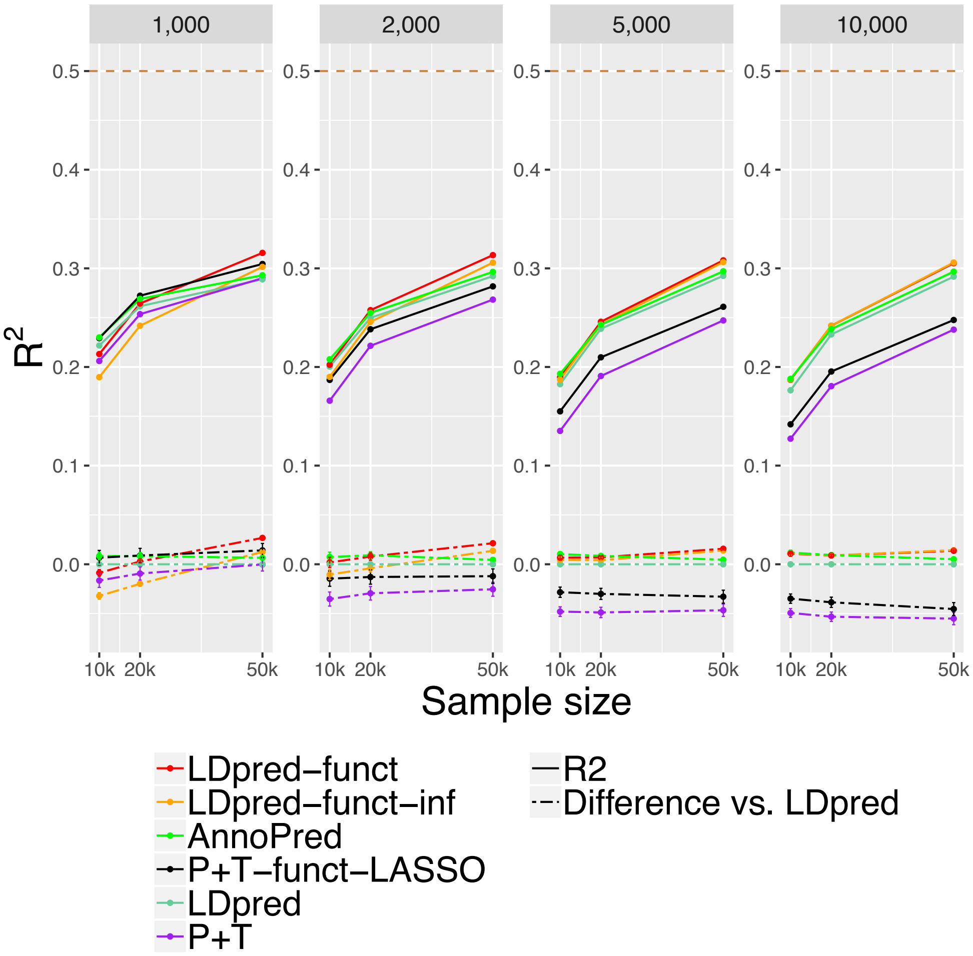


**Figure S1: Accuracy of 6 polygenic prediction methods in simulations using UK Biobank genotypes, for 4 values of the number of causal variants.** We report results for P+T, LDpred, P+T-funct-LASSO, AnnoPred, LDpred-funct-inf and LDpred-funct in chromosome 1 simulations with 1,000 causal variants (extremely sparse architecture), 2,000 causal variants (sparse architecture), 5,000 causal variants (polygenic architecture) and 10,000 causal variants (extremely polygenic architecture). Results are averaged across 100 simulations. Top dashed line denotes simulated SNP-heritability of 0.5. Bottom dashed lines denote differences vs. LDpred; error bars represent 95% confidence intervals. Numerical results are reported in Table S3 and Table S4.

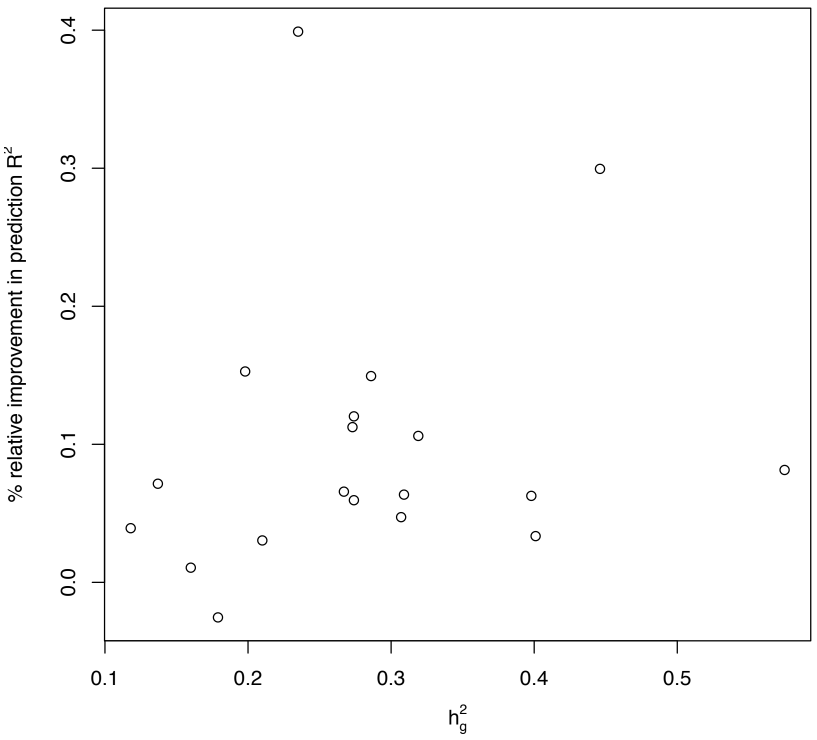


**Figure S2: Relative improvement of LDpred-funct vs. LDpred as a function of $h_g^2$.** We plot relative improvement vs. $h_g^2$ (measured on the observed scale for binary traits) for 19 UK Biobank traits; we excluded two sex-specific traits, age at menarche and balding type I. We observed a correlation of 0.186 across the 19 traits, which was non-significant ($P = 0.4$).

### Supplementary Tables

| | Trait | $h_g^2$ | Training N | Validation N (ancestry distribution) |
|---|---|---|---|---|
| 1 | Height | 0.57 | 408092 | 24351 (40.3% Irish,59.7% Other) |
| 2 | Hair color | 0.45 | 403024 | 24114 (40.3% Irish,59.7% Other) |
| 3 | Platelet count | 0.40 | 395747 | 23616 (40.4% Irish,59.6% Other) |
| 4 | Bone mineral density | 0.40 | 397274 | 23505 (40.4% Irish,59.6% Other) |
| 5 | Red blood cell count | 0.32 | 396464 | 23644 (40.4% Irish,59.6% Other) |
| 6 | Age at menarche | 0.31 | 214860 | 13737 (36.5% Irish,63.5% Other) |
| 7 | FEV1 FVC ratio | 0.31 | 331786 | 19457 (39.5% Irish,60.5% Other) |
| 8 | Body mass index | 0.31 | 407667 | 24322 (40.3% Irish,59.7% Other) |
| 9 | RBC distribution width | 0.29 | 394258 | 23518 (40.3% Irish,59.7% Other) |
| 10 | Forced vital capacity | 0.27 | 331786 | 19457 (39.5% Irish,60.5% Other) |
| 11 | Eosinophil count | 0.27 | 391787 | 23377 (40.3% Irish,59.7% Other) |
| 12 | White blood cell count | 0.27 | 395835 | 23634 (40.4% Irish,59.6% Other) |
| 13 | Systolic Blood pressure | 0.27 | 376437 | 22531 (40.1% Irish,59.9% Other) |
| 14 | Waist hip ratio | 0.21 | 408196 | 24354 (40.3% Irish,59.7% Other) |

**Table S1: List of 14 UK Biobank quantitative traits.** We list the training sample size and validation sample size for each trait. $h_g^2$ estimates are obtained using BOLT-LMM v2.3 using the training data set.

| | Trait | $h_g^2$ | Training | | Validation | |
|---|---|---|---|---|---|---|
| | | | N | Prevalence | N (ancestry distribution) | Prevalence |
| 1 | Balding Type I | 0.32 | 186,506 | 0.32 | 10,171 (45.9% Irish,54.1% Other) | 0.35 |
| 2 | Tanning | 0.23 | 400,721 | 0.39 | 23,947 (40.4% Irish,59.6% Other) | 0.61 |
| 3 | College Education | 0.20 | 405,140 | 0.31 | 24,085 (40.4% Irish,59.6% Other) | 0.50 |
| 4 | Hyperthension | 0.18 | 408,323 | 0.27 | 24,368 (40.3% Irish,59.7% Other) | 0.24 |
| 5 | Cardiovascular Diseases | 0.16 | 408,963 | 0.32 | 24,435 (40.3% Irish,59.7% Other) | 0.29 |
| 6 | Morning Person | 0.14 | 365,245 | 0.37 | 22,188 (40.2% Irish,59.8% Other) | 0.58 |
| 7 | Eczema | 0.12 | 408,454 | 0.23 | 24,374 (40.4% Irish,59.6% Other) | 0.23 |

**Table S2: List of 7 UK Biobank binary traits.** We list the training sample size, validation sample size and prevalence for each trait. $h_g^2$ estimates are obtained using BOLT-LMM v2.3 using the training data set.

| # Causal variants | Model | Training sample size 10,000 Average $R^2(s.e.)$ | 20,000 Average $R^2(s.e.)$ | 50,000 Average $R^2(s.e.)$ |
|---|---|---|---|---|
| 1,000 | P+T | 0.2061 ( 0.0022) | 0.2536 ( 0.0021) | 0.2900 ( 0.0019) |
| | LDpred | 0.2218 ( 0.0024) | 0.2616 ( 0.0021) | 0.2889 ( 0.0018) |
| | P+T-funct-LASSO | 0.2292 ( 0.0024) | 0.2723 ( 0.0024) | 0.3044 ( 0.002) |
| | AnnoPred | 0.2300 ( 0.0025 ) | 0.2691 ( 0.0027 ) | 0.2930 ( 0.0019 ) |
| | LDpred-funct-inf | 0.1896 ( 0.0018) | 0.2419 ( 0.0019) | 0.3015 ( 0.0019) |
| | LDpred-funct | 0.2131 ( 0.002) | 0.2644 ( 0.0021) | 0.3157 ( 0.002) |
| 2,000 | P+T | 0.1658 ( 0.0022) | 0.2215 ( 0.0026) | 0.2683 ( 0.0029) |
| | LDpred | 0.2004 ( 0.0028) | 0.2498 ( 0.0023) | 0.2921 ( 0.0015) |
| | P+T-funct-LASSO | 0.1869 ( 0.0026) | 0.2383 ( 0.0028) | 0.2817 ( 0.0031) |
| | AnnoPred | 0.2078 ( 0.0018 ) | 0.2549 ( 0.0028 ) | 0.2964 ( 0.0016 ) |
| | LDpred-funct-inf | 0.1900 ( 0.0015) | 0.2458 ( 0.0015) | 0.3057 ( 0.0016) |
| | LDpred-funct | 0.2023 ( 0.0016) | 0.2576 ( 0.0016) | 0.3134 ( 0.0017) |
| 5,000 | P+T | 0.1352 ( 0.0016) | 0.1909 ( 0.002) | 0.2472 ( 0.0024) |
| | LDpred | 0.1826 ( 0.0017) | 0.2388 ( 0.0013) | 0.2924 ( 0.0013) |
| | P+T-funct-LASSO | 0.1550 ( 0.0018) | 0.2098 ( 0.0021) | 0.2610 ( 0.0026) |
| | AnnoPred | 0.1931 ( 0.0013 ) | 0.2429 ( 0.0021 ) | 0.2970 ( 0.0014 ) |
| | LDpred-funct-inf | 0.1872 ( 0.0012) | 0.2430 ( 0.0013) | 0.3063 ( 0.0014) |
| | LDpred-funct | 0.1895 ( 0.0012) | 0.2458 ( 0.0013) | 0.3081 ( 0.0014) |
| 10,000 | P+T | 0.1273 ( 0.0015) | 0.1806 ( 0.002) | 0.2379 ( 0.0024) |
| | LDpred | 0.1764 ( 0.0016) | 0.2330 ( 0.0012) | 0.2916 ( 0.0012) |
| | P+T-funct-LASSO | 0.1419 ( 0.0017) | 0.1954 ( 0.0022) | 0.2477 ( 0.0026) |
| | AnnoPred | 0.1880 ( 0.0013 ) | 0.2384 ( 0.0021 ) | 0.2967 ( 0.0013 ) |
| | LDpred-funct-inf | 0.1873 ( 0.0012) | 0.2419 ( 0.0012) | 0.3059 ( 0.0013) |
| | LDpred-funct | 0.1870 ( 0.0013) | 0.2418 ( 0.0012) | 0.3053 ( 0.0012) |

**Table S3: Accuracy of 6 polygenic prediction methods in simulations using UK Biobank genotypes, for 4 values of the number of causal variants.** We report results for P+T, LDpred, P+T-funct-LASSO, AnnoPred, LDpred-funct-inf and LDpred-funct in chromosome 1 simulations with 1,000 causal variants (extremely sparse architecture), 2,000 causal variants (sparse architecture), 5,000 causal variants (polygenic architecture) and 10,000 causal variants (extremely polygenic architecture). Results are averaged across 100 simulations. We report standard errors in parentheses. For SBayesR simulations at N=10K, we obtained average prediction R2 of 0.2754 ( 0.002 ), 0.2381 ( 0.001 ), 0.1983 ( 0.001 ) and 0.1827 ( 0.001) for 1,000, 2,000, 5,000 and 10,000 causal variants respectively. However, for all SBayesR simulations at N=20K and N=50K, we obtained average prediction $R^2$ of 0.0005 (0.0001) and $h_g^2 = 0.99$ in 100/100 simulations, perhaps because the algorithm failed to converge.

**(a)**

| # Causal variants | Model | Training sample size<br>10,000<br>Diff. $R^2 (s.e.)$ | <br>20,000<br>Diff. $R^2 (s.e.)$ | <br>50,000<br>Diff. $R^2 (s.e.)$ |
|---|---|---|---|---|
| 1,000 | P+T | 0.0069 (0.0018) | 0.0106 (0.0016) | 0.0254 (0.0015) |
| | P+T-funct-LASSO | -0.0162 (0.002) | -0.0081 (0.0018) | 0.011 (0.0016) |
| | LDpred | -0.0087 (0.0017) | 0.0028 (0.0013) | 0.0267 (8e-04) |
| | AnnoPred | -0.0169 (0.0018) | -0.0054 (0.0017) | 0.0203 (0.0012) |
| | LDpred-funct-inf | 0.0235 (8e-04) | 0.0225 (6e-04) | 0.0142 (6e-04) |
| | LDpred-funct | 0 | 0 | 0 |
| 2,000 | P+T | 0.0365 (0.0019) | 0.0361 (0.0022) | 0.0451 (0.0026) |
| | P+T-funct-LASSO | 0.0153 (0.0023) | 0.0194 (0.0024) | 0.0317 (0.0027) |
| | LDpred | 0.0019 (0.0026) | 0.0078 (0.0019) | 0.0213 (7e-04) |
| | AnnoPred | -0.0056 (0.0012) | -0.0012 (0.0012) | 0.0167 (7e-04) |
| | LDpred-funct-inf | 0.0123 (5e-04) | 0.0118 (5e-04) | 0.0077 (4e-04) |
| | LDpred-funct | 0 | 0 | 0 |
| 5,000 | P+T | 0.0544 (0.0016) | 0.055 (0.0018) | 0.0609 (0.0021) |
| | P+T-funct-LASSO | 0.0345 (0.0017) | 0.036 (0.0019) | 0.0471 (0.0023) |
| | LDpred | 0.0067 (0.0013) | 0.007 (7e-04) | 0.0157 (5e-04) |
| | AnnoPred | -0.0035 (5e-04) | -0.0015 (6e-04) | 0.011 (6e-04) |
| | LDpred-funct-inf | 0.0023 (3e-04) | 0.0029 (3e-04) | 0.0018 (2e-04) |
| | LDpred-funct | 0 | 0 | 0 |
| 10,000 | P+T | 0.0597 (0.0016) | 0.0612 (0.002) | 0.0674 (0.0024) |
| | P+T-funct-LASSO | 0.0451 (0.0017) | 0.0464 (0.0022) | 0.0576 (0.0026) |
| | LDpred | 0.0107 (0.0013) | 0.0089 (5e-04) | 0.0136 (5e-04) |
| | AnnoPred | -0.0011 (3e-04) | -4e-04 (5e-04) | 0.0085 (5e-04) |
| | LDpred-funct-inf | -4e-04 (2e-04) | -1e-04 (2e-04) | -7e-04 (2e-04) |
| | LDpred-funct | 0 | 0 | 0 |

**(b)**

| # Causal variants | Model | Training sample size<br>10,000<br>Diff. $R^2 (s.e.)$ | <br>20,000<br>Diff. $R^2 (s.e.)$ | <br>50,000<br>Diff. $R^2 (s.e.)$ |
|---|---|---|---|---|
| 1,000 | P+T | -0.0165 (0.0035) | -0.0094 (0.0034) | -2e-04 (0.0033) |
| | LDpred | 0 | 0 | 0 |
| | P+T-funct-LASSO | 0.0067 (0.0037) | 0.0088 (0.0037) | 0.0141 (0.0035) |
| | AnnoPred | 0.0084 (0.0019) | 0.0083 (0.0018) | 0.0066 (0.0011) |
| | LDpred-funct-inf | -0.0321 (0.0017) | -0.0198 (0.0012) | 0.0125 (6e-04) |
| | LDpred-funct | -0.0087 (0.0017) | 0.0028 (0.0013) | 0.0267 (8e-04) |
| 2,000 | P+T | -0.0352 (0.0036) | -0.0294 (0.0035) | -0.0254 (0.0036) |
| | LDpred | 0 | 0 | 0 |
| | P+T-funct-LASSO | -0.0146 (0.0039) | -0.0129 (0.0036) | -0.0121 (0.0037) |
| | AnnoPred | 0.0074 (0.0025) | 0.0091 (0.0019) | 0.0045 (5e-04) |
| | LDpred-funct-inf | -0.0104 (0.0025) | -0.004 (0.0019) | 0.0137 (5e-04) |
| | LDpred-funct | 0.0019 (0.0026) | 0.0078 (0.0019) | 0.0213 (7e-04) |
| 5,000 | P+T | -0.048 (0.0024) | -0.0488 (0.0026) | -0.0466 (0.0031) |
| | LDpred | 0 | 0 | 0 |
| | P+T-funct-LASSO | -0.0283 (0.0026) | -0.03 (0.0028) | -0.0329 (0.0033) |
| | AnnoPred | 0.0103 (0.0012) | 0.0085 (8e-04) | 0.0046 (5e-04) |
| | LDpred-funct-inf | 0.0044 (0.0013) | 0.0041 (7e-04) | 0.0139 (4e-04) |
| | LDpred-funct | 0.0067 (0.0013) | 0.007 (7e-04) | 0.0157 (5e-04) |
| 10,000 | P+T | -0.0493 (0.0022) | -0.0532 (0.0024) | -0.0551 (0.0031) |
| | LDpred | 0 | 0 | 0 |
| | P+T-funct-LASSO | -0.0348 (0.0024) | -0.0386 (0.0026) | -0.0454 (0.0033) |
| | AnnoPred | 0.0118 (0.0013) | 0.0094 (5e-04) | 0.0051 (4e-04) |
| | LDpred-funct-inf | 0.0111 (0.0012) | 0.009 (4e-04) | 0.0143 (5e-04) |
| | LDpred-funct | 0.0107 (0.0013) | 0.0089 (5e-04) | 0.0136 (5e-04) |

**Table S4: Differences between polygenic prediction methods in simulations using UK Biobank genotypes, for 4 values of the number of causal variants.** We report results for P+T, LDpred, P+T-funct-LASSO, AnnoPred, LDpred-funct-inf and LDpred-funct in chromosome 1 simulations with 1,000 causal variants (extremely sparse architecture), 2,000 causal variants (sparse architecture), 5,000 causal variants (polygenic architecture) and 10,000 causal variants (extremely polygenic architecture). Results are averaged across 100 simulations. We report standard errors in parentheses. (a) Difference between $R^2$ for LDpred-funct vs. $R^2$ for each method. (b) Difference between $R^2$ for each method vs. $R^2$ for LDpred.

| | Training sample size | | |
|---|---|---|---|
| # Causal | 10,000 | 20,000 | 50,000 |
| 1,000 | 0.03 | 0.1 | 1 |
| 2,000 | 0.03 | 0.1 | 1 |
| 5,000 | 0.03 | 0.1 | 1 |
| 10,000 | 0.1 | 0.3 | 1 |

**Table S5: Model parameter values for LDpred in simulations.** We report the optimal value of $p$ which is the fraction of non-zero effects in the prior.

| # Causal | | Training sample size 10,000 | 20,000 | 50,000 |
|---|---|---|---|---|
| 1,000 | P+T | 0.0001 | 0.0001 | 0.0001 |
| | P+T-funct-LASSO HP SNP Set | 0.1000 | 0.1000 | 0.3000 |
| | P+T-funct-LASSO LP SNP Set | 0.0100 | 0.0100 | 0.0100 |
| 2,000 | P+T | 0.0010 | 0.0010 | 0.0010 |
| | P+T-funct-LASSO HP SNP Set | 0.1000 | 0.1000 | 0.3000 |
| | P+T-funct-LASSO LP SNP Set | 0.0100 | 0.0100 | 0.0100 |
| 5,000 | P+T | 0.0100 | 0.0100 | 0.0100 |
| | P+T-funct-LASSO HP SNP Set | 0.3000 | 0.3000 | 0.3000 |
| | P+T-funct-LASSO LP SNP Set | 0.1000 | 0.1000 | 0.1000 |
| 10,000 | P+T | 0.1000 | 0.1000 | 0.0100 |
| | P+T-funct-LASSO HP SNP Set | 0.3000 | 0.3000 | 1.0000 |
| | P+T-funct-LASSO LP SNP Set | 0.1000 | 0.1000 | 0.1000 |

**Table S6: Model parameter values for P+T and P+T-funct-LASSO in simulations.** We report the optimal p-value threshold for Pruning + Thresholding (P+T), optimal p-value threshold for P+T-funct-LASSO high prior SNP (HP) set and optimal p-value threshold for P+T-funct-LASSO low prior SNP (LP) set. Optimal $R^2_{LD}$ values was 0.1.

| | Training sample size | | |
|---|---|---|---|
| # Causal | 10,000 | 20,000 | 50,000 |
| 1,000 | 0.03 | 0.10 | 1.00 |
| 2,000 | 0.03 | 0.10 | 1.00 |
| 5,000 | 0.03 | 0.10 | 1.00 |
| 10,000 | 0.10 | 0.10 | 1.00 |

**Table S7: Model parameter values for AnnoPred in simulations.** Model parameter values for AnnoPred in simulations. We report the optimal value of $p$, the proportion of causal variants in the AnnoPred model, for each number of causal variants and sample size simulated.

| Method | Posterior mean effects | LD matrices | Total |
|---|---|---|---|
| P+T | NA | 11 | 11 |
| LDpred | 4,268 (151) | 3734 | 8,003 |
| SBayesR | 142 (9) | 19200 | 19,342 |
| P+T-func-LASSO | NA | 11 | 11 |
| AnnoPred | 5,249 (87) | 3734 | 8,983 |
| LDpred-funct-inf | 71 (10.47) | NA | 71 |
| LDpred-funct | 71 (10.47) | NA | 71 |

**Table S8: Average running time for all 7 methods.** We report average running times in minutes for each method. We separately report the time to estimate posterior mean causal effect sizes, and the time to compute LD matrices (not applicable for LDpred-funct-inf and LD-pred-funct) (we do not include the time to compute polygenic risk scores, which is small in comparison and depends on the number of validation samples). LDpred-funct includes an additional step, the estimation of regularization weights, but this adds minimal additional computational cost (roughly 2 minutes). We estimated running times for chromosome 1 (roughly 10% of the genome) across 20 simulations for estimation of posterior mean effect sizes and one simulation for LD computation, and extrapolated the results to the whole genome. Thus, we report s.e. in parentheses for estimation of posterior mean effect sizes only. LDpred, AnnoPred, LDpred-funct-inf and LDpred-funct require a data coordination step (coordinating the summary statistics file, valditation genotypes and LD reference genotypes (in this case the same as the validation)), which required an additional 193 minutes for LDpred and AnnoPred and 141 minutes for LDpred-funct-inf and LDpred-funct. The running time for computation of the LD matrices for SBayesR is copied from ref. 9 (1.1M SNP set), as we did not repeat this computation.

| # Causal variants | Model | Training sample size: 10,000 Average $R^2 (s.e.)$ | 20,000 Average $R^2 (s.e.)$ | 50,000 Average $R^2 (s.e.)$ |
|---|---|---|---|---|
| 1,000 | P+T | 0.9371 ( 0.0282) | 0.9806 ( 0.0294) | 0.8189 ( 0.0486) |
| | LDpred | 0.992 ( 0.0146 ) | 0.947 ( 0.0083 ) | 0.8521 ( 0.004 ) |
| | P+T-funct-LASSO | 1.5051 ( 0.0589 ) | 1.3703 ( 0.0386 ) | 1.0151 ( 0.075 ) |
| | AnnoPred | 0.8222 ( 0.0171 ) | 0.7832 ( 0.0187 ) | 0.761 ( 0.0154 ) |
| | LDpred-funct-inf | 0.4708 ( 0.0025) | 0.454 ( 0.002) | 0.4345 ( 0.0024) |
| | LDpred-funct | 0.9803 ( 6e-04) | 0.9847 ( 4e-04) | 0.9877 ( 4e-04) |
| 2,000 | P+T | 0.7644 ( 0.0309) | 0.791 ( 0.0257) | 0.7976 ( 0.0209) |
| | LDpred | 0.9688 ( 0.037 ) | 0.9346 ( 0.0257 ) | 0.8483 ( 0.0044 ) |
| | P+T-funct-LASSO | 1.3572 ( 0.0382 ) | 1.2138 ( 0.0544 ) | 1.0448 ( 0.0284 ) |
| | AnnoPred | 0.9258 ( 0.0084 ) | 0.8849 ( 0.0089 ) | 0.8136 ( 0.0025 ) |
| | LDpred-funct-inf | 0.4656 ( 0.004) | 0.457 ( 0.0028) | 0.4396 ( 0.0021) |
| | LDpred-funct | 0.9787 ( 0.001) | 0.9837 ( 7e-04) | 0.9882 ( 4e-04) |
| 5,000 | P+T | 0.4546 ( 0.0207) | 0.5954 ( 0.0172) | 0.6728 ( 0.0158) |
| | LDpred | 0.9984 ( 0.0067 ) | 0.9671 ( 0.0071 ) | 0.8538 ( 0.0044 ) |
| | P+T-funct-LASSO | 0.8085 ( 0.0267 ) | 0.8994 ( 0.012 ) | 0.909 ( 0.0213 ) |
| | AnnoPred | 0.949 ( 0.0034 ) | 0.9092 ( 0.0031 ) | 0.8128 ( 0.0024 ) |
| | LDpred-funct-inf | 0.47 ( 0.0035) | 0.4584 ( 0.0023) | 0.4424 ( 0.0015) |
| | LDpred-funct | 0.9776 ( 9e-04) | 0.9839 ( 5e-04) | 0.9881 ( 4e-04) |
| 10,000 | P+T | 0.3196 ( 0.0136) | 0.4655 ( 0.016) | 0.586 ( 0.0116) |
| | LDpred | 0.9903 ( 0.0156 ) | 0.9449 ( 0.0059 ) | 0.847 ( 0.0041 ) |
| | P+T-funct-LASSO | 0.6824 ( 0.0182 ) | 0.8142 ( 0.0182 ) | 0.8178 ( 0.017 ) |
| | AnnoPred | 0.9468 ( 0.0033 ) | 0.9086 ( 0.0034 ) | 0.811 ( 0.0021 ) |
| | LDpred-funct-inf | 0.4654 ( 0.0028) | 0.4528 ( 0.0025) | 0.4365 ( 0.0024) |
| | LDpred-funct | 0.9761 ( 7e-04) | 0.9824 ( 6e-04) | 0.9874 ( 4e-04) |

**Table S9: Calibration of 6 polygenic prediction methods in simulations using UK Biobank genotypes, for 4 values of the number of causal variants.** We report calibration slopes for P+T, LDpred, P+T-funct-LASSO, AnnoPred, LDpred-funct-inf and LDpred-funct in chromosome 1 simulations with 1,000 causal variants (extremely sparse architecture), 2,000 causal variants (sparse architecture), 5,000 causal variants (polygenic architecture) and 10,000 causal variants (extremely polygenic architecture). Results are averaged across 100 simulations.

| # Causal variants | Model | Training sample size 10,000 | 20,000 | 50,000 |
|---|---|---|---|---|
| | | Average $R^2(s.e.)$ | Average $R^2(s.e.)$ | Average $R^2(s.e.)$ |
| 1,000 | LDpred-funct-inf | 0.1896 ( 0.0018) | 0.2419 ( 0.0019) | 0.3015 ( 0.0019) |
| | LDpred-funct-inf-5 | 0.208 ( 0.002) | 0.2585 ( 0.002) | 0.3104 ( 0.0019) |
| | LDpred-funct-inf-10 | 0.2101 ( 0.002) | 0.261 ( 0.002) | 0.3124 ( 0.002) |
| | LDpred-funct-inf-20 | 0.2116 ( 0.002) | 0.263 ( 0.002) | 0.314 ( 0.002) |
| | LDpred-funct-inf-30 | 0.2126 ( 0.002) | 0.2638 ( 0.002) | 0.315 ( 0.002) |
| | LDpred-funct-inf-40 | 0.2131 ( 0.002) | 0.2644 ( 0.0021) | 0.3157 ( 0.002) |
| | LDpred-funct-inf-50 | 0.2141 ( 0.002) | 0.2652 ( 0.0021) | 0.3161 ( 0.002) |
| | LDpred-funct-inf-60 | 0.2145 ( 0.0021) | 0.2655 ( 0.0021) | 0.3172 ( 0.002) |
| | LDpred-funct-inf-70 | 0.2157 ( 0.0021) | 0.266 ( 0.0021) | 0.317 ( 0.0021) |
| | LDpred-funct-inf-80 | 0.216 ( 0.002) | 0.2665 ( 0.0021) | 0.3173 ( 0.0021) |
| | LDpred-funct-inf-90 | 0.2164 ( 0.0021) | 0.2667 ( 0.0021) | 0.3176 ( 0.0021) |
| | LDpred-funct-inf-100 | 0.2165 ( 0.0021) | 0.267 ( 0.0021) | 0.3174 ( 0.0021) |
| 2,000 | LDpred-funct-inf | 0.1900 ( 0.0015) | 0.2458 ( 0.0015) | 0.3057 ( 0.0016) |
| | LDpred-funct-inf-5 | 0.1994 ( 0.0016) | 0.254 ( 0.0016) | 0.3101 ( 0.0016) |
| | LDpred-funct-inf-10 | 0.2005 ( 0.0016) | 0.2554 ( 0.0016) | 0.3113 ( 0.0017) |
| | LDpred-funct-inf-20 | 0.2016 ( 0.0016) | 0.2566 ( 0.0016) | 0.3124 ( 0.0017) |
| | LDpred-funct-inf-30 | 0.2018 ( 0.0016) | 0.2572 ( 0.0016) | 0.3129 ( 0.0017) |
| | LDpred-funct-inf-40 | 0.2023 ( 0.0016) | 0.2576 ( 0.0016) | 0.3134 ( 0.0017) |
| | LDpred-funct-inf-50 | 0.2023 ( 0.0016) | 0.2575 ( 0.0016) | 0.3136 ( 0.0017) |
| | LDpred-funct-inf-60 | 0.2025 ( 0.0016) | 0.258 ( 0.0017) | 0.3137 ( 0.0017) |
| | LDpred-funct-inf-70 | 0.2027 ( 0.0016) | 0.2579 ( 0.0017) | 0.3135 ( 0.0017) |
| | LDpred-funct-inf-80 | 0.2031 ( 0.0016) | 0.2583 ( 0.0017) | 0.3133 ( 0.0017) |
| | LDpred-funct-inf-90 | 0.2028 ( 0.0016) | 0.2579 ( 0.0017) | 0.3134 ( 0.0018) |
| | LDpred-funct-inf-100 | 0.2031 ( 0.0016) | 0.2582 ( 0.0017) | 0.313 ( 0.0018) |
| 5,000 | LDpred-funct-inf | 0.1872 ( 0.0012) | 0.243 ( 0.0013) | 0.3063 ( 0.0014) |
| | LDpred-funct-inf-5 | 0.1895 ( 0.0012) | 0.2451 ( 0.0013) | 0.3075 ( 0.0014) |
| | LDpred-funct-inf-10 | 0.1898 ( 0.0012) | 0.2456 ( 0.0013) | 0.3079 ( 0.0014) |
| | LDpred-funct-inf-20 | 0.1897 ( 0.0012) | 0.2461 ( 0.0013) | 0.3083 ( 0.0014) |
| | LDpred-funct-inf-30 | 0.1898 ( 0.0012) | 0.2461 ( 0.0013) | 0.3084 ( 0.0014) |
| | LDpred-funct-inf-40 | 0.1895 ( 0.0012) | 0.2458 ( 0.0013) | 0.3081 ( 0.0014) |
| | LDpred-funct-inf-50 | 0.1894 ( 0.0012) | 0.2457 ( 0.0013) | 0.3081 ( 0.0014) |
| | LDpred-funct-inf-60 | 0.1893 ( 0.0012) | 0.2454 ( 0.0013) | 0.3077 ( 0.0014) |
| | LDpred-funct-inf-70 | 0.1891 ( 0.0012) | 0.245 ( 0.0013) | 0.3073 ( 0.0014) |
| | LDpred-funct-inf-80 | 0.1888 ( 0.0012) | 0.2447 ( 0.0013) | 0.3071 ( 0.0014) |
| | LDpred-funct-inf-90 | 0.1885 ( 0.0012) | 0.2444 ( 0.0013) | 0.3066 ( 0.0014) |
| | LDpred-funct-inf-100 | 0.188 ( 0.0012) | 0.244 ( 0.0013) | 0.3062 ( 0.0014) |
| 10,000 | LDpred-funct-inf | 0.1873 ( 0.0012) | 0.2419 ( 0.0012) | 0.3059 ( 0.0013) |
| | LDpred-funct-inf-5 | 0.1883 ( 0.0012) | 0.2428 ( 0.0012) | 0.3064 ( 0.0013) |
| | LDpred-funct-inf-10 | 0.1882 ( 0.0012) | 0.2428 ( 0.0012) | 0.3064 ( 0.0012) |
| | LDpred-funct-inf-20 | 0.1878 ( 0.0012) | 0.2427 ( 0.0012) | 0.3061 ( 0.0012) |
| | LDpred-funct-inf-30 | 0.1873 ( 0.0013) | 0.2422 ( 0.0012) | 0.3056 ( 0.0013) |
| | LDpred-funct-inf-40 | 0.187 ( 0.0013) | 0.2418 ( 0.0012) | 0.3053 ( 0.0012) |
| | LDpred-funct-inf-50 | 0.1865 ( 0.0012) | 0.2414 ( 0.0012) | 0.3049 ( 0.0013) |
| | LDpred-funct-inf-60 | 0.186 ( 0.0013) | 0.2409 ( 0.0012) | 0.3043 ( 0.0013) |
| | LDpred-funct-inf-70 | 0.1855 ( 0.0013) | 0.2406 ( 0.0012) | 0.3039 ( 0.0013) |
| | LDpred-funct-inf-80 | 0.1851 ( 0.0012) | 0.2399 ( 0.0012) | 0.3036 ( 0.0012) |
| | LDpred-funct-inf-90 | 0.1846 ( 0.0013) | 0.2393 ( 0.0012) | 0.3027 ( 0.0013) |
| | LDpred-funct-inf-100 | 0.1841 ( 0.0013) | 0.2387 ( 0.0012) | 0.3027 ( 0.0013) |

**Table S10: Sensitivity of LDpred-funct results to number of bins used for regularization in simulations using UK Biobank genotypes**. We report results with the number of posterior mean causal effect size bins used for regularization ($K$) set to 10, 20, 50 or 100. LDpred-funct-$K$ denotes each respective value of $K$. We also report results for LDpred-funct-inf, which is identical to LDpred-funct with $K$ set to 1. Results are averaged across 100 simulations. We report standard errors in parentheses.

| | | Training sample size | | |
|---|---|---|---|---|
| # Causal | | 10,000 | 20,000 | 50,000 |
| variants | Model | Average $R^2(s.e.)$ | Average $R^2(s.e.)$ | Average $R^2(s.e.)$ |
| 1,000 | LDpred-funct-inf | 0.1896 ( 0.0018) | 0.2419 ( 0.0019) | 0.3015 ( 0.0019) |
| | LDpred-funct | 0.2131 ( 0.002) | 0.2644 ( 0.0021) | 0.3157 ( 0.002) |
| | LDpred-funct-inf-cheat | 0.1926 ( 0.0018) | 0.2456 ( 0.0019) | 0.3074 ( 0.002) |
| | LDpred-funct-cheat | 0.2221 ( 0.0021) | 0.2714 ( 0.0022) | 0.3228 ( 0.0021) |
| 2,000 | LDpred-funct-inf | 0.1900 ( 0.0015) | 0.2458 ( 0.0015) | 0.3057 ( 0.0016) |
| | LDpred-funct | 0.2023 ( 0.0016) | 0.2576 ( 0.0016) | 0.3134 ( 0.0017) |
| | LDpred-funct-inf-cheat | 0.1943 ( 0.0015) | 0.2498 ( 0.0016) | 0.3108 ( 0.0016) |
| | LDpred-funct-cheat | 0.2109 ( 0.0016) | 0.2646 ( 0.0017) | 0.3193 ( 0.0017) |
| 5,000 | LDpred-funct-inf | 0.1872 ( 0.0012) | 0.243 ( 0.0013) | 0.3063 ( 0.0014) |
| | LDpred-funct | 0.1895 ( 0.0012) | 0.2458 ( 0.0013) | 0.3081 ( 0.0014) |
| | LDpred-funct-inf-cheat | 0.1928 ( 0.0013) | 0.2479 ( 0.0013) | 0.3102 ( 0.0014) |
| | LDpred-funct-cheat | 0.1972 ( 0.0014) | 0.252 ( 0.0013) | 0.3121 ( 0.0014) |
| 10,000 | LDpred-funct-inf | 0.1873 ( 0.0012) | 0.2419 ( 0.0012) | 0.3059 ( 0.0013) |
| | LDpred-funct | 0.1870 ( 0.0013) | 0.2418 ( 0.0012) | 0.3053 ( 0.0012) |
| | LDpred-funct-inf-cheat | 0.1937 ( 0.0012) | 0.2474 ( 0.0012) | 0.3097 ( 0.0012) |
| | LDpred-funct-cheat | 0.194 ( 0.0013) | 0.2482 ( 0.0013) | 0.3096 ( 0.0013) |

**Table S11: Accuracy of LDpred-funct method in simulations using UK Biobank genotypes under different BaselineLD estimates, for 4 values of the number of causal variants.** LDpred-funct-cheat refers to a ”cheating” version of LDpred-funct that utilized the true baseline-LD model parameters used to simulate the data. Results are averaged across 100 simulations.

| # Causal variants | Model | Training sample size 10,000 Average $R^2 (s.e.)$ | 20,000 Average $R^2 (s.e.)$ | 50,000 Average $R^2 (s.e.)$ |
|---|---|---|---|---|
| 2,000 | P+T | 0.0634 ( 0.0015 ) | 0.0912 ( 0.0019 ) | 0.118 ( 0.0056 ) |
| | LDpred | 0.0885 ( 0.0025 ) | 0.1174 ( 0.0029 ) | 0.1447 ( 0.0028 ) |
| | P+T-funct-LASSO | 0.0695 ( 0.0022 ) | 0.0988 ( 0.0021 ) | 0.1213 ( 0.006 ) |
| | AnnoPred | 0.0848 ( 0.0024 ) | 0.1097 ( 0.0021 ) | 0.134 ( 0.0023 ) |
| | LDpred-funct-inf | 0.0759 ( 0.0019 ) | 0.1033 ( 0.0021 ) | 0.1384 ( 0.0023 ) |
| | LDpred-funct | 0.0808 ( 0.0023 ) | 0.1094 ( 0.0023 ) | 0.1432 ( 0.0022 ) |
| 5,000 | P+T | 0.0503 ( 0.0013 ) | 0.0783 ( 0.0014 ) | 0.1033 ( 0.0051 ) |
| | LDpred | 0.0746 ( 0.0016 ) | 0.1041 ( 0.0016 ) | 0.1382 ( 0.0021 ) |
| | P+T-funct-LASSO | 0.0578 ( 0.0015 ) | 0.0869 ( 0.0015 ) | 0.1087 ( 0.0056 ) |
| | AnnoPred | 0.0772 ( 0.0016 ) | 0.1031 ( 0.0025 ) | 0.1334 ( 0.0019 ) |
| | LDpred-funct-inf | 0.0744 ( 0.0014 ) | 0.1032 ( 0.0016 ) | 0.1375 ( 0.0018 ) |
| | LDpred-funct | 0.0754 ( 0.0015 ) | 0.105 ( 0.0017 ) | 0.139 ( 0.0018 ) |

**Table S12: Accuracy of 6 polygenic prediction methods in simulations with lower SNP-heritability ($h_g^2 = 0.25$) using UK Biobank genotypes, for 4 values of the number of causal variants.** We report results for P+T, LDpred, P+T-funct-LASSO, AnnoPred, LDpred-funct-inf and LDpred-funct in chromosome 1 simulations with 1,000 causal variants (extremely sparse architecture), 2,000 causal variants (sparse architecture), 5,000 causal variants (polygenic architecture) and 10,000 causal variants (extremely polygenic architecture). Results are averaged across 20 simulations. We report standard errors in parentheses. For SBayesR simulations at N=10K, we obtained average prediction $R^2$ of 0.0625 ( 0.0086 ) and 0.0618 ( 0.0053 ) for 2,000, and 5,000 causal variants respectively. However, for all SBayesR simulations at N=20K and N=50K, we obtained average prediction $R^2$ of 0.0002 (0.0001) and $h_g^2 = 0.99$ in 20/20 simulations, perhaps because the algorithm failed to converge.

| | Trait | Training $N$ | $h_g^2$ | $c$ | bins |
|---|---|---|---|---|---|
| 1 | Height | 408092 | 0.57 | 0.45 | 100 |
| 2 | Hair color | 403024 | 0.45 | 0.22 | 100 |
| 3 | Platelet count | 395747 | 0.40 | 0.29 | 88 |
| 4 | Bone mineral density | 397274 | 0.40 | 0.26 | 87 |
| 5 | Red blood cell count | 396464 | 0.32 | 0.21 | 70 |
| 6 | Age at menarche | 214860 | 0.31 | 0.20 | 40 |
| 7 | FEV1 FVC ratio | 331786 | 0.31 | 0.24 | 56 |
| 8 | Body mass index | 407667 | 0.31 | 0.27 | 70 |
| 9 | RBC distribution width | 394258 | 0.29 | 0.20 | 63 |
| 10 | Eosinophil count | 391787 | 0.27 | 0.18 | 60 |
| 11 | Forced vital capacity | 331786 | 0.27 | 0.22 | 50 |
| 12 | White blood cell count | 395835 | 0.27 | 0.21 | 60 |
| 13 | Systolic Blood pressure | 376437 | 0.27 | 0.21 | 56 |
| 14 | Waist hip ratio | 408196 | 0.21 | 0.15 | 48 |
| 1 | Balding type I | 186506 | 0.32 | 0.11 | 31 |
| 2 | Tanning ability | 400721 | 0.23 | 0.09 | 53 |
| 3 | College Education | 405140 | 0.20 | 0.15 | 45 |
| 4 | Hyperthension | 408323 | 0.18 | 0.14 | 41 |
| 5 | Cardiovascular Diseases | 408963 | 0.16 | 0.12 | 37 |
| 6 | Morning Person | 365245 | 0.14 | 0.11 | 29 |
| 7 | Eczema | 408454 | 0.12 | 0.09 | 27 |

**Table S13: Parameter values for 21 UK Biobank traits.** The 14 quantitative traits are listed first, followed by the 7 binary traits. For each trait, we list the training sample size, $h_g^2$ estimate (from BOLT-LMM v2.3; used by LDpred, LDpred-funct-inf and LDpred-funct), the $c$ parameter (used by LDpred-funct-inf and LDpred-funct) and number of bins for LDpred-funct.

| | Trait | $h_g^2$ | P+T | LDpred | SBayesR | P+T-funct-LASSO | AnnoPred | LDpred -funct-inf | LDpred -funct |
|---|---|---|---|---|---|---|---|---|---|
| 1 | Height | 0.575 | 0.3481 (0.0145) | 0.3820 (0.0204) | 0.3820 (0.0197) | 0.3684 (0.0172) | 0.4063 (0.0276) | 0.4003 (0.0196) | 0.4131 (0.0261) |
| 2 | Hair color | 0.446 | 0.2418 (0.0794) | 0.2507 (0.1024) | 0.3241 (0.1236) | 0.2413 (0.0874) | 0.2616 (0.1095) | 0.2638 (0.1059) | 0.3258 (0.1305) |
| 3 | Platelet count | 0.401 | 0.2044 (0.0228) | 0.2349 (0.0227) | 0.2338 (0.0264) | 0.2187 (0.0245) | 0.2333 (0.0221) | 0.2290 (0.0202) | 0.2428 (0.0275) |
| 4 | Bone mineral density | 0.398 | 0.1906 (0.0190) | 0.2096 (0.0216) | 0.2176 (0.0193) | 0.2024 (0.0206) | 0.2281 (0.0210) | 0.2107 (0.0186) | 0.2227 (0.0248) |
| 5 | Red blood cell count | 0.319 | 0.1244 (0.0143) | 0.1494 (0.0158) | 0.1523 (0.0167) | 0.1309 (0.0149) | 0.1676 (0.0158) | 0.1568 (0.0141) | 0.1652 (0.0207) |
| 6 | Age at menarche | 0.313 | 0.0716 (0.0054) | 0.1078 (0.0100) | 0.1057 (0.0098) | 0.0873 (0.0066) | 0.1113 (0.0103) | 0.1054 (0.0090) | 0.1078 (0.0185) |
| 7 | FEV1 FVC ratio | 0.309 | 0.1012 (0.0077) | 0.1238 (0.0095) | 0.1348 (0.0098) | 0.1124 (0.0076) | 0.1417 (0.0099) | 0.1289 (0.0090) | 0.1317 (0.0172) |
| 8 | Body mass index | 0.307 | 0.1071 (0.0044) | 0.1414 (0.0076) | 0.1402 (0.0072) | 0.1186 (0.0059) | 0.1519 (0.0079) | 0.1491 (0.0073) | 0.1481 (0.0152) |
| 9 | RBC distribution width | 0.286 | 0.1190 (0.0141) | 0.1302 (0.0147) | 0.1399 (0.0157) | 0.1299 (0.0177) | 0.1458 (0.0147) | 0.1394 (0.0145) | 0.1496 (0.0203) |
| 10 | Eosinophil count | 0.274 | 0.1109 (0.0102) | 0.1259 (0.0133) | 0.1288 (0.0131) | 0.1169 (0.0102) | 0.1391 (0.0144) | 0.1342 (0.0130) | 0.1410 (0.0194) |
| 11 | Forced vital capacity | 0.274 | 0.0763 (0.0054) | 0.1083 (0.0072) | 0.1070 (0.0069) | 0.0893 (0.0053) | 0.1196 (0.0074) | 0.1153 (0.0068) | 0.1147 (0.0152) |
| 12 | White blood cell count | 0.273 | 0.0964 (0.0069) | 0.1145 (0.0095) | 0.1256 (0.0095) | 0.1085 (0.0093) | 0.1325 (0.0094) | 0.1244 (0.0090) | 0.1274 (0.0160) |
| 13 | Systolic Blood pressure | 0.267 | 0.0827 (0.0055) | 0.1066 (0.0071) | 0.1084 (0.0069) | 0.0967 (0.0053) | 0.1195 (0.0071) | 0.1135 (0.0066) | 0.1136 (0.0138) |
| 14 | Waist hip ratio | 0.210 | 0.0567 (0.0039) | 0.0782 (0.0062) | 0.0782 (0.0064) | 0.0646 (0.0046) | 0.0855 (0.0069) | 0.0786 (0.0051) | 0.0806 (0.0117) |
| 15 | Average | 0.330 | 0.1379 (0.002) | 0.1617 (0.0098) | 0.1699 (0.0108) | 0.1490 (0.008) | 0.1745 (0.0103) | 0.1678 (0.0098) | 0.1774 (0.0113) |

**Table S14: Accuracy of 7 polygenic prediction methods across 14 UK Biobank quantitative traits.** We report results for P+T, LDpred, SBayesR, P+T-funct-LASSO, AnnoPred, LDpred-funct-inf and LDpred-funct. Optimal parameters for each method are reported in Table S19, Table S18, Table S21 and Table S13. We report block jackknife standard error over 200 equally sized blocks of adjacent SNPs.

| | Trait | $h_g^2$ | P+T | LDpred | SBayesR | P+T-funct-LASSO | AnnoPred | LDpred -funct-inf | LDpred -funct |
|---|---|---|---|---|---|---|---|---|---|
| 1 | Balding type I | 0.323 | 0.1164 (0.0129) | 0.1326 (0.0171) | 0.1272 (0.0197) | 0.1272 (0.0162) | 0.1461 (0.0211) | 0.1078 (0.0135) | 0.1237 (0.0245) |
| 2 | Tanning ability | 0.235 | 0.1293 (0.0445) | 0.1278 (0.0670) | 0.1688 (0.0678) | 0.1298 (0.0467) | 0.1199 (0.0633) | 0.1219 (0.0626) | 0.1788 (0.0771) |
| 3 | College Education | 0.198 | 0.0611 (0.0028) | 0.0687 (0.0063) | 0.0711 (0.0061) | 0.0636 (0.0029) | 0.0694 (0.0064) | 0.0770 (0.0061) | 0.0792 (0.0113) |
| 4 | Hyperthension | 0.179 | 0.0405 (0.0028) | 0.0537 (0.0046) | 0.0575 (0.0044) | 0.0467 (0.0032) | 0.0553 (0.0047) | 0.0527 (0.0044) | 0.0523 (0.0093) |
| 5 | Cardiovascular Diseases | 0.160 | 0.0275 (0.0020) | 0.0424 (0.0038) | 0.0464 (0.0039) | 0.0335 (0.0025) | 0.0451 (0.0039) | 0.0432 (0.0037) | 0.0429 (0.0085) |
| 6 | Morning Person | 0.137 | 0.0265 (0.0021) | 0.0382 (0.0033) | 0.0382 (0.0031) | 0.0308 (0.0025) | 0.0395 (0.0032) | 0.0413 (0.0032) | 0.0409 (0.0087) |
| 7 | Eczema | 0.118 | 0.0149 (0.0016) | 0.0273 (0.0032) | 0.0254 (0.0029) | 0.0210 (0.0025) | 0.0307 (0.0034) | 0.0279 (0.0031) | 0.0284 (0.0070) |
| 8 | Average | 0.190 | 0.0594 (0.0015) | 0.0701 (0.0103) | 0.0764 (0.0103) | 0.0647 (0.0073) | 0.0723 (0.0100) | 0.0674 (0.0095) | 0.0780 (0.0116) |

**Table S15: Accuracy of 7 polygenic prediction methods across 7 UK Biobank binary traits.** We report results for P+T, LDpred, SBayesR, P+T-funct-LASSO, AnnoPred, LDpred-funct-inf and LDpred-funct. Optimal parameters for each method are reported in Table S19, Table S18, Table S21 and Table S13. We report block jackknife standard error over 200 equally sized blocks of adjacent SNPs.

| Phenotype | Average difference vs LDpred | Average difference vs LDpred-funct |
|---|---|---|
| P+T | -0.0194 (0.0081) | -0.0325 (0.0094) |
| LDPRED | 0.000 (0.0000) | -0.0131 (0.0034) |
| SBayesR | 0.0067 (0.0031) | -0.0064 (0.0031) |
| P+T-LASSO | -0.0108 (0.003) | -0.0239 (0.004) |
| Annopred | 0.0093 (0.002) | -0.0038 (0.0034) |
| LDPRED-funct-inf | 0.0032 (0.0019) | -0.0099 (0.0026) |
| LDPRED-funct | 0.0131 (0.0034) | 0.000 (0.0000) |

**Table S16: Average absolute differences between polygenic prediction methods across 21 UK Biobank traits.** We report results for P+T, LDpred, SBayesR, P+T-funct-LASSO, AnnoPred, LDpred-funct-inf and LDpred-funct. We report the difference between prediction $R^2$ for each method vs. prediction $R^2$ for LDpred, and difference between prediction $R^2$ for each method vs. prediction $R^2$ for LDpred-funct. Block-jackknife standard errors on the differences are reported in parentheses.

| | Method | Average $R^2$ |
|---|---|---|
| 1 | P+T | 0.1112 |
| 2 | LDpred | 0.1311 |
| 3 | SBayesR | 0.1379 |
| 4 | P+T-funct-LASSO | 0.1203 |
| 5 | AnnoPred | 0.1405 |
| 6 | LDpred-funct-inf | 0.1343 |
| 7 | LDpred-funct | 0.1443 |
| 8 | LDpred-inf | 0.1126 |
| 9 | LDpred (without excluding long-range LD regions) | 0.0839 |
| 10 | LDpred (typed SNPs only) | 0.1299 |
| 11 | LDpred-funct-inf (typed SNPs only) | 0.1124 |
| 12 | LDpred-funct (typed SNPs only) | 0.1198 |
| 13 | SBayesR (2.9M) | 0.1227 |
| 14 | P+T-funct-LASSO-weighted | 0.1231 |
| 15 | P+T-funct-LASSO (5%) | 0.1219 |
| 16 | LDpred-funct-inf (meta31) | 0.1307 |
| 17 | P+T-LASSO (random) | 0.1178 |
| 18 | AnnoPred (random) | 0.1288 |
| 19 | LDpred-funct-inf (random) | 0.1153 |
| 20 | LDpred-funct (random) | 0.1271 |
| 21 | LDpred-inf + sparsity | 0.1282 |
| 22 | LDpred-funct-inf (baseline) | 0.1312 |
| 23 | LDpred-funct (baseline) | 0.1415 |
| 24 | LDpred-funct-inf (constant prior) | 0.1241 |
| 25 | LDpred-funct (constant prior) | 0.1383 |
| 26 | LDpred-funct-inf(QCfilters) | 0.1339 |
| 27 | LDpred-funct-inf(UK10K) | 0.1354 |
| 28 | LDpred-funct-inf(UK10K, baseline-LD+LDAK) | 0.1350 |
| 29 | LDpred-funct-inf (Baseline-LD v2.1) | 0.1360 |
| 30 | LDpred-funct (Baseline-LD v2.1) | 0.1469 |

**Table S17: Accuracy of secondary polygenic prediction methods across 21 UK Biobank traits.** Rows 1-7 correspond to average prediction $R^2$ across 21 UK Biobank traits for each method. Row 8 correspond to the average prediction $R^2$ from LDpred-inf. Row 9 correspond to the average prediction $R^2$ from LDpred that includes SNPs from long-range LD regions. Rows 10-12 are methods that analyze only genotyped SNPs (601,728 genotyped SNPs after QC). Row 13 correspond to the average prediction $R^2$ from SBayesR using 2.9M SNPs (SNP set described in Methods). Rows 14-15 are slightly modified versions of P+T-funct-LASSO. Row 16 uses baseline-LD model functional enrichments that were meta-analyzed across 31 traits. Rows 17-20 report. avg. prediction $R^2$ functionally informed methods, P+T-funct-LASSO, AnnoPred, LDpred-funct-inf and LDpred-funct using a set of 75 random annotations. Row 21 corresponds to avg. prediction $R^2$ of LDpred-inf with sparsity (i.e. LDpred-funct with no functional annotations except the annotation containing all SNPs). Row 22-23 uses the baseline model, instead of the baseline-LD model. Rows 24-25 report avg. prediction $R^2$ for LDpred-funct-inf and LDpred-funct with priors replaced by indicator function that gives a constant prior to SNPs with predicted $h^2 > 0$ and sets effect sizes to zero for SNPs with $h^2 < 0$. Row 26 restricts the baseline-LD model to the 6,334,603 SNPs that passed QC filters and were used for prediction. Row 27 infers baseline-LD model parameters using UK10K SNPs, instead of 1000 Genomes SNPs. Row 28 uses UK10K SNPs and uses the baseline-LD+LDAK model, instead of the baseline-LD model. Row 29-30 corresponds to the average prediction $R^2$ for LDpred-funct-inf and LDpred-funct using baseline-LD model v2.1 (instead of baseline-LD model v1.1, which is used in our main analyses).

| | Trait | $h_g^2$ | $p$ |
|---|---|---|---|
| 1 | Height | 0.57 | 0.3000 |
| 2 | Hair color | 0.45 | 0.3000 |
| 3 | Platelet count | 0.40 | 0.1000 |
| 4 | Bone mineral density | 0.40 | 0.1000 |
| 5 | Balding type I | 0.32 | 0.0300 |
| 6 | Red blood cell count | 0.32 | 0.1000 |
| 7 | Age at menarche | 0.31 | 0.0300 |
| 8 | FEV1 FVC ratio | 0.31 | 0.1000 |
| 9 | Body mass index | 0.31 | 0.1000 |
| 10 | RBC distribution width | 0.29 | 0.1000 |
| 11 | Forced vital capacity | 0.27 | 0.0300 |
| 12 | Eosinophil count | 0.27 | 0.0300 |
| 13 | White blood cell count | 0.27 | 0.1000 |
| 14 | Systolic Blood pressure | 0.27 | 0.1000 |
| 15 | Tanning ability | 0.23 | 0.1000 |
| 16 | Waist hip ratio | 0.21 | 0.0300 |
| 17 | College Education | 0.20 | 0.1000 |
| 18 | Hyperthension | 0.18 | 0.0300 |
| 19 | Cardiovascular Diseases | 0.16 | 0.0300 |
| 20 | Morning Person | 0.14 | 0.0100 |
| 21 | Eczema | 0.12 | 0.0100 |

**Table S18: Model parameter values for LDpred applied to 21 UK Biobank traits.** $h_g^2$ estimate (from BOLT-LMM v2.3), $p$ is the fraction of non-zero effects in the prior.

| | Phenotype | $h_g^2$ | P+T | P-values threshold for P+T-funct-LASSO HP SNP set | P+T-funct-LASSO LP SNP set |
|---|---|---|---|---|---|
| 1 | Height | 0.57 | 0.0100 | 0.30 | 0.10 |
| 2 | Hair color | 0.45 | 0.0010 | 0.30 | 0.01 |
| 3 | Platelet count | 0.40 | 0.0010 | 0.10 | 0.10 |
| 4 | Bone mineral density | 0.40 | 0.0010 | 0.10 | 0.10 |
| 5 | Balding type I | 0.32 | 0.0010 | 0.10 | 0.01 |
| 6 | Red blood cell count | 0.32 | 0.0010 | 0.10 | 0.10 |
| 7 | Age at menarche | 0.31 | 0.0100 | 0.10 | 0.10 |
| 8 | FEV1 FVC ratio | 0.31 | 0.0010 | 0.10 | 0.10 |
| 9 | Body mass index | 0.31 | 0.1000 | 0.30 | 0.10 |
| 10 | RBC distribution width | 0.29 | 0.0010 | 0.10 | 0.01 |
| 11 | Forced vital capacity | 0.27 | 0.0010 | 0.10 | 0.10 |
| 12 | Eosinophil count | 0.27 | 0.0010 | 0.10 | 0.10 |
| 13 | White blood cell count | 0.27 | 0.0100 | 0.10 | 0.10 |
| 14 | Systolic Blood pressure | 0.27 | 0.0010 | 0.10 | 0.10 |
| 15 | Tanning ability | 0.23 | 0.0001 | 0.10 | 0.01 |
| 16 | Waist hip ratio | 0.21 | 0.0100 | 0.10 | 0.10 |
| 17 | College Education | 0.20 | 1.0000 | 0.30 | 0.30 |
| 18 | Hyperthension | 0.18 | 0.0100 | 0.10 | 0.01 |
| 19 | Cardiovascular Diseases | 0.16 | 0.0010 | 0.10 | 0.01 |
| 20 | Morning Person | 0.14 | 0.0100 | 0.10 | 0.10 |
| 21 | Eczema | 0.12 | 0.0001 | 0.10 | 0.01 |

**Table S19: Model parameter values for P+T and P+T-funct-LASSO in 21 UK Biobank traits.** We report the optimal p-value threshold for Pruning + Thresholding (P+T), optimal p-value threshold for P+T-funct-LASSO high prior SNP (HP) set and optimal p-value threshold for P+T-funct-LASSO low prior SNP (LP) set. Optimal $R^2_{LD}$ values was 0.1.

| | Phenotype | $h_g^2$ | Absolute difference |
|---|---|---|---|
| 1 | Height | 0.575 | 0.0343 (0.0048) |
| 2 | Hair color | 0.446 | 0.0238 (0.0136) |
| 3 | Platelet count | 0.401 | 0.0352 (0.0045) |
| 4 | Bone mineral density | 0.398 | 0.0268 (0.0031) |
| 5 | Balding type I | 0.323 | 0.0230 (0.0031) |
| 6 | Red blood cell count | 0.319 | 0.0325 (0.0042) |
| 7 | Age at menarche | 0.313 | 0.0151 (0.0027) |
| 8 | FEV1 FVC ratio | 0.309 | 0.0202 (0.0030) |
| 9 | Body mass index | 0.307 | 0.0126 (0.0019) |
| 10 | RBC distribution width | 0.286 | 0.0388 (0.0074) |
| 11 | Forced vital capacity | 0.274 | 0.0198 (0.0021) |
| 12 | Eosinophil count | 0.274 | 0.0388 (0.0062) |
| 13 | White blood cell count | 0.273 | 0.0239 (0.0037) |
| 14 | Systolic Blood pressure | 0.267 | 0.0155 (0.0021) |
| 15 | Tanning ability | 0.235 | 0.0394 (0.0183) |
| 16 | Waist hip ratio | 0.210 | 0.0142 (0.0018) |
| 17 | College Education | 0.198 | 0.0052 (0.0028) |
| 18 | Hyperthension | 0.179 | 0.0074 (0.0013) |
| 19 | Cardiovascular Diseases | 0.160 | 0.0060 (0.0012) |
| 20 | Morning Person | 0.137 | 0.0042 (0.0015) |
| 21 | Eczema | 0.118 | 0.0073 (0.0010) |
| 22 | Average across traits | 0.286 | 0.0211 (0.0020) |

**Table S20: Difference between LDpred-funct-inf and LDpred-inf.** LDpred-funct-inf significantly outperforms LDpred-inf ($P < 10^{-20}$ for difference using one-sided z-test based on block-jackknife standard error)

| | Trait | $h_g^2$ | $p$ |
|---|---|---|---|
| 1 | Height | 0.57 | 0.3000 |
| 2 | Hair color | 0.45 | 1.0000 |
| 3 | Platelet count | 0.40 | 0.3000 |
| 4 | Bone mineral density | 0.40 | 0.3000 |
| 5 | Balding type I | 0.32 | 0.0300 |
| 6 | Red blood cell count | 0.32 | 0.3000 |
| 7 | Age at menarche | 0.31 | 0.1000 |
| 8 | FEV1 FVC ratio | 0.31 | 0.1000 |
| 9 | Body mass index | 0.31 | 0.1000 |
| 10 | RBC distribution width | 0.29 | 0.3000 |
| 11 | Forced vital capacity | 0.27 | 0.1000 |
| 12 | Eosinophil count | 0.27 | 0.1000 |
| 13 | White blood cell count | 0.27 | 0.1000 |
| 14 | Systolic Blood pressure | 0.27 | 0.3000 |
| 15 | Tanning ability | 0.23 | 0.3000 |
| 16 | Waist hip ratio | 0.21 | 0.1000 |
| 17 | College Education | 0.20 | 0.3000 |
| 18 | Hyperthension | 0.18 | 0.1000 |
| 19 | Cardiovascular Diseases | 0.16 | 0.1000 |
| 20 | Morning Person | 0.14 | 0.1000 |
| 21 | Eczema | 0.12 | 0.1000 |

**Table S21: Model parameter values for AnnoPred applied to 21 UK Biobank traits.** We report the $h_g^2$ estimate (from BOLT-LMM v2.3) and the optimal value of $p$, the proportion of causal variants in the AnnoPred model.

| | Trait | $h_g^2$ | LDpred-funct-inf | Validation sample size 1000 | 2000 | 5000 | 10000 | ALL |
|---|---|---|---|---|---|---|---|---|
| 1 | Height | 0.57 | 0.4003 | 0.3970 | 0.4048 | 0.4108 | 0.4122 | 0.4131 |
| 2 | Hair color | 0.45 | 0.2638 | 0.2957 | 0.3010 | 0.3052 | 0.3081 | 0.3258 |
| 3 | Platelet count | 0.40 | 0.2290 | 0.2469 | 0.2425 | 0.2451 | 0.2432 | 0.2428 |
| 4 | Bone mineral density | 0.40 | 0.2107 | 0.2350 | 0.2300 | 0.2274 | 0.2242 | 0.2227 |
| 5 | Balding type I | 0.32 | 0.1078 | 0.1280 | 0.1227 | 0.1233 | 0.1233 | 0.1237 |
| 6 | Red blood cell count | 0.32 | 0.1568 | 0.1692 | 0.1643 | 0.1678 | 0.1642 | 0.1652 |
| 7 | Age at menarche | 0.31 | 0.1054 | 0.1143 | 0.1128 | 0.1121 | 0.1114 | 0.1078 |
| 8 | FEV1 FVC ratio | 0.31 | 0.1289 | 0.1365 | 0.1357 | 0.1327 | 0.1348 | 0.1317 |
| 9 | Body mass index | 0.31 | 0.1491 | 0.1530 | 0.1477 | 0.1522 | 0.1520 | 0.1481 |
| 10 | RBC distribution width | 0.29 | 0.1394 | 0.1513 | 0.1458 | 0.1528 | 0.1546 | 0.1496 |
| 11 | Forced vital capacity | 0.27 | 0.1153 | 0.1265 | 0.1165 | 0.1118 | 0.1123 | 0.1147 |
| 12 | Eosinophil count | 0.27 | 0.1342 | 0.1464 | 0.1415 | 0.1416 | 0.1432 | 0.1410 |
| 13 | White blood cell count | 0.27 | 0.1244 | 0.1325 | 0.1280 | 0.1284 | 0.1288 | 0.1274 |
| 14 | Systolic Blood pressure | 0.27 | 0.1135 | 0.1200 | 0.1109 | 0.1141 | 0.1109 | 0.1136 |
| 15 | Tanning ability | 0.23 | 0.1219 | 0.1521 | 0.1582 | 0.1786 | 0.1796 | 0.1788 |
| 16 | Waist hip ratio | 0.21 | 0.0786 | 0.0919 | 0.0842 | 0.0822 | 0.0801 | 0.0806 |
| 17 | College Education | 0.20 | 0.0770 | 0.0775 | 0.0771 | 0.0739 | 0.0717 | 0.0792 |
| 18 | Hyperthension | 0.18 | 0.0527 | 0.0554 | 0.0572 | 0.0542 | 0.0528 | 0.0523 |
| 19 | Cardiovascular Diseases | 0.16 | 0.0432 | 0.0494 | 0.0486 | 0.0432 | 0.0428 | 0.0429 |
| 20 | Morning Person | 0.14 | 0.0413 | 0.0417 | 0.0427 | 0.0383 | 0.0366 | 0.0409 |
| 21 | Eczema | 0.12 | 0.0279 | 0.0335 | 0.0313 | 0.0290 | 0.0276 | 0.0284 |
| 22 | Average across traits | 0.29 | 0.1343 | 0.1454 | 0.1430 | 0.1440 | 0.1435 | 0.1443 |

**Table S22: Sensitivity of LDpred-funct results to number of validation samples across 21 UK Biobank traits.** We report results with the number of validation samples set to 1,000, 2,000, 5,000, 10,000 (the number of regularization bins is proportional to the number of validation samples; see Equation 6. Results are averaged across 100 random subsets of each size. ALL denotes results of LDpred-funct using the total number of validation samples (reported in Table S1). We also report results for LDpred-funct-inf, which is equivalent to LDpred-funct in the limit of a very small number of validation samples.

| | Phenotype | $h_g^2$ | LDpred-funct-inf | LDpred-funct | LDpred-funct (1K training weights) |
|---|---|---|---|---|---|
| 1 | Height | 0.57 | 0.4003 | 0.4131 | 0.4130 |
| 2 | Hair color | 0.45 | 0.2638 | 0.3258 | 0.3258 |
| 3 | Platelet count | 0.40 | 0.2290 | 0.2428 | 0.2427 |
| 4 | Bone mineral density | 0.40 | 0.2107 | 0.2227 | 0.2226 |
| 5 | Balding type I | 0.32 | 0.1078 | 0.1237 | 0.1238 |
| 6 | Red blood cell count | 0.32 | 0.1568 | 0.1652 | 0.1653 |
| 7 | Age at menarche | 0.31 | 0.1054 | 0.1078 | 0.1080 |
| 8 | FEV1 FVC ratio | 0.31 | 0.1289 | 0.1317 | 0.1317 |
| 9 | Body mass index | 0.31 | 0.1491 | 0.1481 | 0.1480 |
| 10 | RBC distribution width | 0.29 | 0.1394 | 0.1496 | 0.1497 |
| 11 | Forced vital capacity | 0.27 | 0.1153 | 0.1147 | 0.1147 |
| 12 | Eosinophil count | 0.27 | 0.1342 | 0.1410 | 0.1410 |
| 13 | White blood cell count | 0.27 | 0.1244 | 0.1274 | 0.1274 |
| 14 | Systolic Blood pressure | 0.27 | 0.1135 | 0.1136 | 0.1136 |
| 15 | Tanning ability | 0.23 | 0.1219 | 0.1788 | 0.1787 |
| 16 | Waist hip ratio | 0.21 | 0.0786 | 0.0806 | 0.0806 |
| 17 | College Education | 0.20 | 0.0770 | 0.0792 | 0.0793 |
| 18 | Hyperthension | 0.18 | 0.0527 | 0.0523 | 0.0524 |
| 19 | Cardiovascular Diseases | 0.16 | 0.0432 | 0.0429 | 0.0429 |
| 20 | Morning Person | 0.14 | 0.0413 | 0.0409 | 0.0410 |
| 21 | Eczema | 0.12 | 0.0279 | 0.0284 | 0.0284 |
| 22 | Average across traits | 0.29 | 0.1343 | 0.1443 | 0.1443 |

**Table S23: Sensitivity of LDpred-funct results restricted to only 1,000 samples to estimate $\alpha_k$ weights across 21 UK Biobank traits.** We report results with the number of validation samples set to 1,000, 2,000, 5,000, 10,000 (the number of regularization bins is proportional to the number of validation samples; see Equation 6. Results are averaged across 100 random subsets of each size. ALL denotes results of LDpred-funct using the total number of validation samples (reported in Table S1). We also report results for LDpred-funct-inf, which is equivalent to LDpred-funct in the limit of a very small number of validation samples.

| | Trait | $h_g^2$ | P+T | LDpred | SBayesR | P+T-funct-LASSO | AnnoPred | LDpred -funct-inf | LDpred -funct |
|---|---|---|---|---|---|---|---|---|---|
| 1 | Height | 0.575 | 0.2228 | 0.7595 | 1.0335 | 0.3034 | 0.7374 | 0.7367 | 0.9938 |
| 2 | Hair color | 0.446 | 0.2505 | 0.7254 | 0.9909 | 0.3058 | 0.7118 | 0.7182 | 0.9920 |
| 3 | Platelet count | 0.401 | 0.2429 | 0.8451 | 0.9645 | 0.3423 | 0.7678 | 0.8115 | 0.9895 |
| 4 | Bone mineral density | 0.398 | 0.2871 | 0.8192 | 0.9604 | 0.3477 | 0.8142 | 0.8246 | 0.9865 |
| 5 | Balding type I | 0.323 | 0.3693 | 0.8994 | 1.9855 | 0.5050 | 0.8034 | 0.8781 | 0.9776 |
| 6 | Red blood cell count | 0.319 | 0.2898 | 0.8583 | 0.9721 | 0.3458 | 0.8481 | 0.8202 | 0.9822 |
| 7 | Age at menarche | 0.313 | 0.1990 | 1.0227 | 0.6183 | 0.3430 | 0.8880 | 0.8706 | 0.9782 |
| 8 | FEV1 FVC ratio | 0.309 | 0.3021 | 0.8843 | 1.0014 | 0.3593 | 0.8591 | 0.8527 | 0.9740 |
| 9 | Body mass index | 0.307 | 0.1687 | 0.9138 | 1.0338 | 0.3541 | 0.8839 | 0.8599 | 0.9813 |
| 10 | RBC distribution width | 0.286 | 0.2839 | 0.8399 | 0.9454 | 0.4189 | 0.8287 | 0.8123 | 0.9833 |
| 11 | Forced vital capacity | 0.274 | 0.2237 | 0.9085 | 1.0890 | 0.3783 | 0.9106 | 0.8665 | 0.9770 |
| 12 | Eosinophil count | 0.274 | 0.2781 | 0.9082 | 0.9874 | 0.3298 | 0.8167 | 0.8518 | 0.9830 |
| 13 | White blood cell count | 0.273 | 0.2352 | 0.9033 | 1.0254 | 0.3707 | 0.8347 | 0.8538 | 0.9793 |
| 14 | Systolic Blood pressure | 0.267 | 0.2200 | 0.9050 | 1.0063 | 0.3637 | 0.9064 | 0.8453 | 0.9808 |
| 15 | Tanning ability | 0.235 | 0.2437 | 0.8312 | 1.8516 | 0.2873 | 0.7384 | 0.8292 | 0.9905 |
| 16 | Waist hip ratio | 0.210 | 0.2057 | 0.8453 | 0.9891 | 0.3344 | 0.8404 | 0.8500 | 0.9758 |
| 17 | College Education | 0.198 | 0.1345 | 1.0159 | 2.2891 | 0.2610 | 0.8747 | 0.8520 | 0.9728 |
| 18 | Hyperthension | 0.179 | 0.2140 | 0.9817 | 2.1853 | 0.3557 | 0.8328 | 0.8077 | 0.9710 |
| 19 | Cardiovascular Diseases | 0.160 | 0.1213 | 0.9376 | 2.1111 | 0.3296 | 0.8329 | 0.7953 | 0.9643 |
| 20 | Morning Person | 0.137 | 0.2158 | 1.0803 | 2.1999 | 0.3720 | 0.9299 | 0.8751 | 0.9651 |
| 21 | Eczema | 0.118 | 0.1752 | 0.7496 | 2.1296 | 0.4971 | 0.8271 | 0.7611 | 0.9634 |
| 22 | Average across traits | 0.286 | 0.2325 | 0.8873 | 1.3509 | 0.3574 | 0.8327 | 0.8273 | 0.9791 |

**Table S24: Calibration comparison for the 7 methods applied to 21 UK Biobank traits.** We report calibration slopes for each method, where a value close to 1 respresents a well calibrated prediction.

| | Trait | LDpred-funct-inf | LDpred-funct-10 | LDpred-funct-20 | LDpred-funct-50 | LDpred-funct-75 | LDpred-funct-100 |
|---|---|---|---|---|---|---|---|
| 1 | Height | 0.4003 | 0.4113 | 0.4116 | 0.4126 | 0.4127 | 0.4128 |
| 2 | Hair color | 0.2624 | 0.2998 | 0.3059 | 0.3174 | 0.3199 | 0.3290 |
| 3 | Platelet count | 0.2315 | 0.2445 | 0.2453 | 0.2445 | 0.2446 | 0.2448 |
| 4 | Bone mineral density | 0.2137 | 0.2266 | 0.2266 | 0.2271 | 0.2265 | 0.2256 |
| 5 | Balding type I | 0.1075 | 0.1217 | 0.1235 | 0.1220 | 0.1198 | 0.1185 |
| 6 | Red blood cell count | 0.1571 | 0.1651 | 0.1655 | 0.1660 | 0.1660 | 0.1649 |
| 7 | Age at menarche | 0.1082 | 0.1118 | 0.1116 | 0.1122 | 0.1112 | 0.1070 |
| 8 | FEV1 FVC ratio | 0.1311 | 0.1353 | 0.1348 | 0.1343 | 0.1336 | 0.1315 |
| 9 | Body mass index | 0.1508 | 0.1501 | 0.1504 | 0.1494 | 0.1481 | 0.1473 |
| 10 | RBC distribution width | 0.1421 | 0.1527 | 0.1535 | 0.1535 | 0.1530 | 0.1517 |
| 11 | Forced vital capacity | 0.1145 | 0.1160 | 0.1155 | 0.1145 | 0.1128 | 0.1118 |
| 12 | Eosinophil count | 0.1335 | 0.1425 | 0.1422 | 0.1415 | 0.1406 | 0.1395 |
| 13 | White blood cell count | 0.1239 | 0.1278 | 0.1284 | 0.1276 | 0.1266 | 0.1261 |
| 14 | Systolic Blood pressure | 0.1114 | 0.1129 | 0.1119 | 0.1118 | 0.1108 | 0.1105 |
| 15 | Tanning ability | 0.1229 | 0.1716 | 0.1794 | 0.1818 | 0.1873 | 0.1892 |
| 16 | Waist hip ratio | 0.0793 | 0.0818 | 0.0810 | 0.0804 | 0.0798 | 0.0782 |
| 17 | College Education | 0.0716 | 0.0720 | 0.0731 | 0.0731 | 0.0748 | 0.0739 |
| 18 | Hyperthension | 0.0523 | 0.0542 | 0.0541 | 0.0528 | 0.0521 | 0.0519 |
| 19 | Cardiovascular Diseases | 0.0423 | 0.0437 | 0.0433 | 0.0421 | 0.0410 | 0.0410 |
| 20 | Morning Person | 0.0372 | 0.0372 | 0.0366 | 0.0359 | 0.0349 | 0.0340 |
| 21 | Eczema | 0.0274 | 0.0278 | 0.0275 | 0.0271 | 0.0274 | 0.0258 |
| 22 | Average across traits | 0.1343 | 0.1432 | 0.1439 | 0.1442 | 0.1440 | 0.1436 |

**Table S25: Sensitivity of LDpred-funct results to number of bins used for regularization across 21 UK Biobank traits.** We report results with the number of posterior mean causal effect size bins used for regularization ($K$) set to 10, 20, 50, 75 or 100. LDpred-funct-$K$ denotes each respective value of $K$. We also report results for LDpred-funct-inf, which is identical to LDpred-funct with $K$ set to 1.

| Data Set | Training $N$ | P+T | LDpred | P+T-funct -LASSO | AnnoPred | LDpred-funct-inf | LDpred-funct |
|---|---|---|---|---|---|---|---|
| UK Biobank interim release | 113660 | 0.2289 | 0.2265 | 0.2609 | 0.2530 | 0.2465 | 0.2506 |
| UK Biobank | 408092 | 0.3448 | 0.3823 | 0.3644 | 0.4063 | 0.4010 | 0.4152 |
| 23andMe | 698430 | 0.2997 | 0.2953 | 0.3141 | 0.3166 | 0.3203 | 0.3442 |
| Meta-analysis of UK Biobank and 23andMe | 1107430 | 0.3710 | 0.3990 | 0.3778 | 0.4189 | 0.4193 | 0.4310 |
| Fixed-effect meta-analysis | 1107430 | 0.3687 | 0.3688 | 0.3663 | 0.3982 | 0.3991 | 0.4142 |

**Table S26: Accuracy of 6 prediction methods in height meta-analysis of UK Biobank and 23andMe cohorts.** We report results for P+T, LDpred, P+T-funct-LASSO, AnnoPred, LDpred-funct-inf and LDpred-funct, for each of 4 training data sets: UK Biobank interim release (113,660 training samples), UK Biobank (408,092 training samples), 23andMe (698,430 training samples) and meta-analysis of UK Biobank and 23andMe (1,107,430 training samples). We also report results for a fixed-effect meta-analysis of UK Biobank and 23andMe. For SBayesR we obtained prediction $R^2$ of 0.0219, 0.3829, 0.2658, 0.3967 and 0.3028 respectively for the 5 data sets listed. For the UK Biobank interim release analysis, we obtained prediction $R^2$ of 0.0219 and $h_g^2 = 0.054$, perhaps because the algorithm failed to converge.

| Method | South Asians | Africans |
|---|---|---|
| Method | Avg $R^2$ | Avg $R^2$ |
| P+T | 0.0568 (0.0022) | 0.0172 (0.0014) |
| LDpred | 0.0666 (0.0031) | 0.0241 (0.0018 ) |
| SBayesR | 0.0724 (0.0032) | 0.0250 (0.0019) |
| P+T-funct-LASSO | 0.0631(0.0025) | 0.0185 (0.0016) |
| AnnoPred | 0.0744 (0.0032) | 0.0282 (0.0036) |
| LDpred-funct-inf | 0.0704 (0.0029) | 0.0263 (0.0019) |
| LDpred-funct | 0.0727 (0.0030) | 0.0296 (0.0019) |

**Table S27: Average prediction accuracy of 7 polygenic prediction methods in individuals of South Asian and African ancestry across 21 UK Biobank traits applied.** We report the average prediction $R^2$ of each method in each population (block-jackknife s.e. in parentheses). We analyzed training samples of British ancestry from UK Biobank (average N=373K). We analyzed 7,444 unrelated South Asian validation samples from UK Biobank (74% Indian, 23% Pakistani, 3% Bangladeshi) and 7,379 unrelated African validation samples from UK Biobank (55% Caribbean, 45% African). In the South Asian validation sample, LDpred-funct attained +9.2%, +0.5%, -2.3%, +3.3% relative improvements in average prediction $R^2$ vs. LDpred, SBayesR, AnnoPred, LDpred-funct-inf respectively; $P < 5 * 10^{-5}$, $P = 0.8$, $P = 0.34$, $P < 7 * 10^{-5}$ for differences using two-sided z-test based on block-jackknife s.e. In the African validation sample, LDpred-funct attained +23%, +18%, +5.0%, +13% relative improvements in average prediction $R^2$ vs. LDpred, SBayesR, AnnoPred, LDpred-funct-inf respectively; $P < 10^{-5}$, $P < 2 * 10^{-3}$, $P = 0.2$, $P < 2 * 10^{-5}$ for differences using two-sided z-test based on block-jackknife s.e.

| | Trait | $h_g^2$ | LDpred-funct-inf under different priors: baselineLD (1000G) | baselineLD (UK10K) | baselineLD + LDAK (UK10K) |
|---|---|---|---|---|---|
| 1 | Height | 0.575 | 0.4003 | 0.4022 | 0.4046 |
| 2 | Hair color | 0.446 | 0.2638 | 0.2778 | 0.2727 |
| 3 | Platelet count | 0.401 | 0.2290 | 0.2291 | 0.2303 |
| 4 | Bone mineral density | 0.398 | 0.2107 | 0.2109 | 0.2108 |
| 5 | Balding type I | 0.323 | 0.1078 | 0.1065 | 0.1082 |
| 6 | Red blood cell count | 0.319 | 0.1568 | 0.1571 | 0.1551 |
| 7 | Age at menarche | 0.313 | 0.1054 | 0.1052 | 0.1051 |
| 8 | FEV1 FVC ratio | 0.309 | 0.1289 | 0.1284 | 0.1286 |
| 9 | Body mass index | 0.307 | 0.1491 | 0.1498 | 0.1484 |
| 10 | RBC distribution width | 0.286 | 0.1394 | 0.1408 | 0.1404 |
| 11 | Eosinophil count | 0.274 | 0.1342 | 0.1350 | 0.1348 |
| 12 | Forced vital capacity | 0.274 | 0.1153 | 0.1150 | 0.1139 |
| 13 | White blood cell count | 0.273 | 0.1244 | 0.1254 | 0.1249 |
| 14 | Systolic Blood pressure | 0.267 | 0.1135 | 0.1144 | 0.1141 |
| 15 | Tanning ability | 0.235 | 0.1219 | 0.1227 | 0.1206 |
| 16 | Waist hip ratio | 0.210 | 0.0786 | 0.0786 | 0.0785 |
| 17 | College Education | 0.198 | 0.0770 | 0.0780 | 0.0777 |
| 18 | Hyperthension | 0.179 | 0.0527 | 0.0533 | 0.0536 |
| 19 | Cardiovascular Diseases | 0.160 | 0.0432 | 0.0434 | 0.0433 |
| 20 | Morning Person | 0.137 | 0.0413 | 0.0414 | 0.0415 |
| 21 | Eczema | 0.118 | 0.0279 | 0.0289 | 0.0282 |

**Table S28: Accuracy of LDpred-funct-inf(1000G), LDpred-funct-inf(UK10K) and LDpred-funct-inf(UK10K, baseline-LD+LDAK) across 21 UK Biobank traits.** We report results for each trait. Results for Average across traits are reported in Table S17.
